# Evaluation of immunotoxic effects and metabolism of enniatins and beauvericin using *in vitro* models of the human intestinal epithelium

**DOI:** 10.64898/2026.10.30.763082

**Authors:** Anita Solhaug, Lada Ivanova, Gunnar Sundstøl Eriksen, Christiane Kruse Fæste

**Author notes:** Corresponding author: Anita Solhaug, Norwegian Veterinary Institute, P.O. Box 64, 1431 Ås, Norway.

## Abstract

Enniatins (ENNs) and beauvericin (BEA) are mycotoxins produced by *Fusarium* species. They are frequently detected in cereal-based foods such as maize and wheat. Consequently, the gastrointestinal tract is a primary site of exposure for consumers of grain-containing foods. This study examined intestinal effects of four major ENNs: ENNB, ENNB1, ENNA, and ENNA1, and the structurally related mycotoxin BEA, using the primary human intestinal model EpiIntestinal™ tissues and a Caco-2/TH29-MTX-E12 co-culture. None of the compounds induced cytotoxicity or impaired the barrier integrity at concentrations ranging from 1 to 10 µM. Furthermore, all ENNs and BEA stimulated IL-6 secretion, with ENNA and ENNA1 inducing the strongest responses, while IL-8 secretion was only triggered by ENNA and ENNA1. Transport studies revealed that approximately 32% of ENNB and 25% of ENNB1 crossed from the apical to the basolateral side of the intestinal tissue within 4 h, whereas the other ENNs and BEA had transport ratios below 10%. Further findings suggested that uptake and intracellular accumulation of ENNs and BEA were influenced by their affinity to the efflux transporter P-glycoprotein (P-gp). In addition, several ENNB and ENNB1 metabolites were generated in the EpiIntestinal™ tissues and transported. Lastly, it was demonstrated that ENNB can disturb the metabolism of the sex hormone estradiol in human liver microsomes. Overall, ENNs and BEA showed limited acute toxicity in human intestinal models but elicited distinct inflammatory responses and differed in transport and metabolism. These findings may be of relevance when evaluating health risks from chronic, low-level dietary exposure.

## 1. Introduction

Enniatins (ENNs) and beauvericin (BEA) are fungal metabolites produced by several species of *Fusarium* fungi. These toxins are closely related cyclic hexadepsipeptides consisting of alternating D-hydroxy-isovaleryl-(2-hydroxy-3-methylbutanoic acid) and N-methylphenylalanine residues (**Fig. 1**), which are highly lipophilic and can form ion channels in biological membranes (EFSA 2014). ENNs occur worldwide, most commonly in cereals, but also in other crop plants (Coulet et al. 2025; EFSA 2014; Mihalache et al. 2023). Despite the widespread occurrence, the knowledge about their toxicity is still limited. As a consequence, EFSA has not yet changed its 2014 decision that the lack of relevant toxicity and toxicokinetic data does not allow assessment of risks to human health and the establishment of tolerable daily intakes (TDI) for ENNs and BEA (EFSA 2014). Recently, an extensive review of existing toxicological knowledge on this mycotoxin group identified data gaps, especially in immunotoxicity and toxicokinetics, including biotransformation and compound interactions (Behr et al. 2025). Most available toxicity studies focus on ENNB, while data on other ENNs and BEA are scarce. *In vivo* immunomodulatory effects of BEA and ENNs have been reported in Atlantic salmon (Søderstrøm et al. 2023) and mice (Behr et al. 2025), but the transferability and relevance of these results to humans must be verified in species-specific models (Leist and Hartung 2013). In this respect, experimental designs that reproduce conditions in the human intestine are particularly important, since foodborne mycotoxins can directly affect the gut epithelium during uptake.

**Fig. 1:**
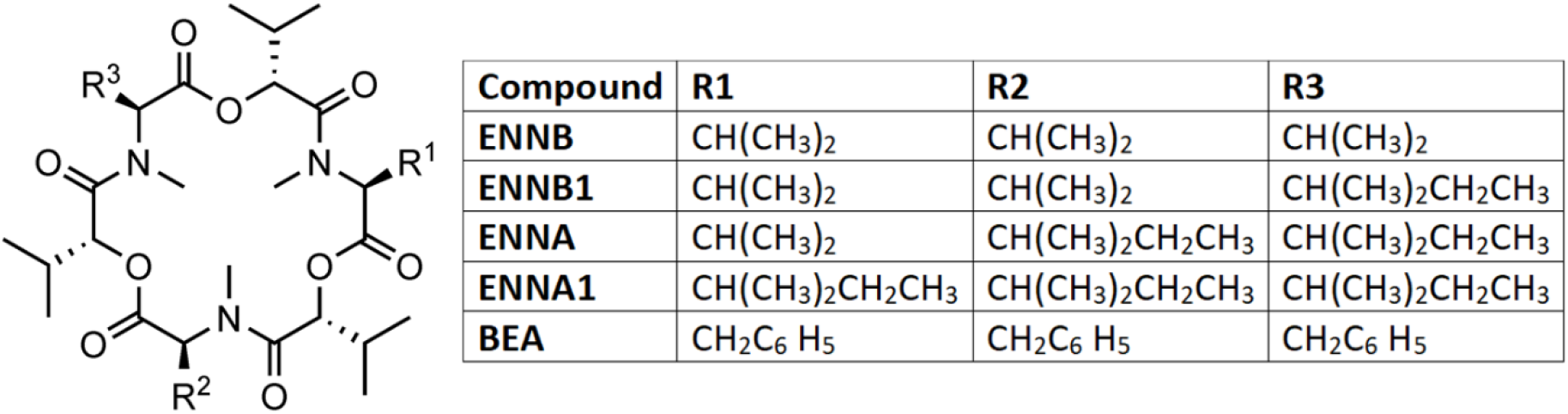
Structure formula of ENNs and BEA (Created in ChemDrawPrime, version 23.0.1.10).

The gastrointestinal tract acts as a selective barrier against dietary contaminants and pathogens while allowing nutrient and water absorption. This physical and chemical barrier consists of an outer mucus layer and a continuous monolayer of epithelial cells joined by tight junctions, adherens junctions, and desmosomes (Vancamelbeke and Vermeire 2017). The underlying lamina propria contains immune cells that support defence mechanisms essential for maintaining mucosal immunity (Neurath et al. 2025). In a healthy gut, physiologically important substances are absorbed or actively transported across the epithelial layer from the apical (AP) lumen to the basolateral (BL) side and into the bloodstream. At the same time, the mucosal immune system tolerates the high load of antigens in gut contents and protects against external pathogens, e.g., through cytokine production. However, disruption of the intestinal barrier by toxic substances or infections can increase the permeability of the gut and lead to a proinflammatory immune response that can progress into a chronic inflammatory disorder or other gut-related diseases (Neurath et al. 2025). The suggested immunomodulatory potential of ENNs and BEA is therefore highly relevant to the assessment because they are widely used in basic food products and, consequently, are commonly found in gut contents.

*In vitro* models commonly used to identify potential intestinal toxicity of ENNs and BEA include the human intestinal epithelial cell lines Caco-2, HT-29, HCT15 and HCT116 (Behr et al. 2025; Elesh et al. 2026; Ranaldi et al. 2026), the porcine cell line IPEC-J2 (Novak et al. 2019), and the rainbow trout cell line RTgut-GC (Dayeh et al. 2025). These studies have predominantly explored the cytotoxicity, trans-membrane transport, and ionophoric properties of these mycotoxins, finding that levels above 10 µM were generally needed to see a significant increase in markers of cell damage. Moreover, ENNs were shown to be substrates of the multidrug resistance protein P-glycoprotein (P-gp) (Dornetshuber et al. 2009; Hiraga et al. 2005; Ivanova et al. 2010). In contrast, only a few experiments have described a possible impact of non-cytotoxic ENNs and BEA levels on the intestinal immune system, as reflected in small changes in proinflammatory cytokine expression (Elesh et al. 2026; Ranaldi et al. 2026).

The Caco-2 cell line, derived from a human colorectal carcinoma, is a well-characterised and widely used cell culture system that, when differentiated under specific culture conditions, displays enterocyte-like morphology and functions (Engle et al. 1998). However, Caco-2 cells contain only negligible levels of the cytochrome P450 enzymes CYP3A4, CYP2C9, and CYP2C19 responsible for xenobiotic biotransformation in the small intestine epithelium (Küblbeck et al. 2016). Furthermore, the variable expression of several transmembrane transporters, lack of mucus production, and overexpression of tight junctions, makeing the cell layer too tight for normal paracellular transport, reduces the suitability of this cell model for the prediction of intestinal substance absorption (Fedi et al. 2021; Markus et al. 2021). However, genetic modifications of some Caco-2 clones have improved this model by enabling expression of CYP enzymes or the formation of an apical brush border (Elesh et al. 2026; Küblbeck et al. 2016). In addition, co-culturing with mucus-producing cell lines such as HT29-MTX-E12, a goblet cell-like clone of human colorectal adenocarcinoma HT-29 cells, in transwell inserts results in a cell monolayer with improved tight junction functionality and an intact mucus layer, resembling the *in vivo* situation (Behrens et al. 2001; Béduneau et al. 2014; Pan et al. 2015).

A more recent approach to optimise absorption models is using organotypic 3D small intestine tissues made from primary human cells, including enterocytes, Paneth cells, M cells, tuft cells, and intestinal stem cells, provided by Mattek Corporation as EpiIntestinal™ tissues. This model has a more *in vivo-*like physiology because it consists of several layers with correct epithelial polarity, mimicking the structure and functionality of the human small intestine. The epithelial layer forms villi and tight junctions as well as expresses brush border enzymes, drug-metabolising enzymes and transport proteins. The EpiIntestinal™ Full Thickness (EpiIntestinal™FT) tissues also include the underlying lamina propria containing human intestinal fibroblasts, making the model useful for studies of inflammation and innate immune responses, since fibroblasts initiate the immune response to tissue injuries by presenting immune cell receptors, inducing the release of cytokines (Cui et al. 2020; Fedi et al. 2021; Markus et al. 2021; Robinson et al. 2023). Multiple pharmaceutical studies have demonstrated the physiological relevance of EpiIntestinal™ tissues for assessing intestinal permeability, drug absorption, enterocyte metabolism, and tissue responses (Ayehunie et al. 2018; Cui et al. 2020; Markus et al. 2021). However, this model remains largely unexplored for studying human *in vivo* exposure to dietary mycotoxins and is considered a New Approach Methodology that could help reduce animal testing in risk assessment (Snapkow et al. 2024).

Transport across the epithelium and biotransformation in enterocytes are key factors determining the absorption kinetics of compounds from the gut lumen into the bloodstream. Depending on the effectiveness of compound-specific transport mechanisms and the degree of metabolism, external and systemic exposure can differ considerably, affecting bioavailability and, in turn, expected efficacy, toxicity, and associated health risks. Available human absorption data for ENNs and BEA, including intestinal biotransformation, are very limited or non-existent (Behr et al. 2025). Bioavailability values from *in vivo* animal studies vary widely; for example, for ENNB1 they range from 5% in chicken to 91% in pig (Devreese et al. 2014; Fraeyman et al. 2016), leaving significant uncertainty about human absorption and systemic exposure. Furthermore, ENNs are extensively metabolised as reported in *in vitro* studies using hepatic microsomes of animals and humans, showing significant differences between species in the metabolic profiles (Fæste et al. 2011; Ivanova et al. 2011). Information on BEA metabolism is even scarcer. Hepatic metabolites have been determined *in vitro* for several species (Elesh et al. 2026; Yuan et al. 2022), but data on absorption or intestinal biotransformation in humans are completely lacking (Behr et al. 2025).

So far, studies exploring the toxicity of ENNs and BEA have all reported negligible acute toxicity after dietary intake, whereas there is increasing evidence for chronic toxicity from frequent low-level oral exposure (Behr et al. 2025). Beyond the growing evidence of immune effects, some indications suggest that cyclic hexadepsipeptidic mycotoxins could disrupt physiological pathways such as steroidogenesis (Kalayou et al. 2015; Lori et al. 2026). In human adrenal gland cells, exposure to 10 µM ENNB reduced hormone concentrations and affected the expression of genes involved in hormone regulation (Kalayou et al. 2015). Furthermore, ENNB1, ENNA1 and BEA demonstrated endocrine-disrupting potential by acting as antagonists of human estrogen and androgen receptors (Jeong et al. 2024; Park and Lee 2021). ENNs and BEA can also directly interact with important endogenous metabolites, for example, during biotransformation or transport processes. Previously, ENNB1 was shown to inhibit CYP 3A-dependent oxidation reactions of typical substrates such as midazolam, but also to be susceptible itself to changes in metabolic efficiency by the presence of other xenobiotics such as the mycotoxin deoxynivalenol (Ivanova et al. 2019). A metabolic interaction between ENNs/BEA and endogenous sex hormones such as estradiol (E2) could thus contribute to the mycotoxins’ effects on hormone homeostasis.

This study aimed to address some of these knowledge gaps by using Caco-2/HT29-MTX-E12 co-cultures and EpiIntestinal™ tissues to examine inflammatory responses and uptake mechanisms of ENNB, ENNB1, ENNA, ENNA1 and BEA in the human gastrointestinal tract. Moreover, the generation and distribution of ENNB- and ENNB1-metabolites and possible interactions with the metabolism of the endogenous sex hormone E2 were evaluated.

## 2. Materials and Methods

### 2.1. Toxins and exposure experiments

ENNB, ENNB1, ENNA and ENNA1 were synthesised by F. Reith, L. Oltmanns, V. Prisyazhnoy and R.D. Süssmuth (Technical University of Berlin, Germany) according to Hu et al. (2012). Ester building blocks were prepared from modified and asymmetrically protected N-methylated amino acids and fused by Ghosez amide coupling. After cyclisation the product was purified by crystallisation. The identity of the respective mycotoxin was confirmed by analytical HPLC-ESI-MS/MS and ^1^HNMR spectroscopy. Beauvericin (BEA, #1142621129) was purchased from Cayman Chemical (Ann Arbor, MI, USA). The ENNs and BEA were dissolved in dimethylsulphoxide (DMSO) or methanol (MeOH) at 5 or 10 mM and stored in glass vials in aliquots at −20 °C. MeOH stocks were used for experiments that involved LC-MSMS analysis, while DMSO stocks were used for all other experiments. Because of the high lipophilicity of ENNs and BEA (Tran et al. 2020), the exposure medium, containing 1 to 30 µM was prepared in glass vials to minimise attachment to plastic surfaces (Solhaug et al. 2025). Estradiol (E2) was purchased from Sigma-Aldrich (Burlington, MA, USA) and diluted in MeOH to a stock concentration of 3.67 mM. Upon exposure, the cell culture medium was replaced with exposure medium containing the respective toxin. The final DMSO or MeOH concentration was 0.5%, and all exposures within the same experiment had similar DMSO/MeOH concentrations.

### 2.2. Cell cultivation

The Caco-2 (#09042001) and HT29-MTX-E12 (#12040401) cell lines were obtained from the European Collection of Authenticated Cell Cultures (ECACC, Porton Down, Salisbury, UK). The cell lines were cultivated in separate 72 cm^2^ vented flasks with Dulbecco’s Modified Eagle medium (DMEM glutamax, Gibco, Thermo Fisher Scientific, Waltham, MA, USA) supplemented with 1% non-essential amino acids (NEAA, Gibco), 1 % penicillin-streptomycin (Gibco), and 10% heat-inactivated foetal bovine serum (FBS, USA origin, Gibco) at 37 °C, 5% CO_2_. The cell lines were routinely screened for mycoplasma contamination using the MycoAlert^®^ Mycoplasma Detection Kit (Lonza, Basel, Switzerland). Subcultivation was performed twice a week, when the culture reached approximately 70% confluence, using TrypLE (Gibco). *Caco-2/HT29-MTX-E12 co-culture*: The cells (9:1 ratio) were plated into hanging cell culture transwell inserts (Millicell^®^ cell culture inserts, 0.4 µm, #PTHT24H48, Sigma-Aldrich) at a density of 1 × 10^5^ cells/cm^2^. After 21 days, with medium refreshed every Monday, Wednesday, and Friday, they were used for cytotoxicity, barrier, and MUC5AC and cytokine secretion experiments. *Caco-2 monoculture:* Caco-2 cells were seeded into standing cell culture transwell inserts (Millicell^®^ cell culture inserts, 0.4 µm, 0.6 cm^2^, #PIHP01250, Sigma-Aldrich) at a density of 1 × 10^5^ cells/cm^2^. The cells were grown for 21 days, with medium refreshed every Monday, Wednesday, and Friday, and used in transport experiments. *Primary human intestinal cells:* EpiIntestinal^TM^ tissues (provided pre-seeded in standing inserts, 24-well format, 0.6 cm^2^) were obtained from Mattek (Sartorius, Bratislava, Slovakia) and cultured according to the manufacturer’s instructions using the provided cell culture medium. The EpiIntestinal^TM^ (#SMI-100) model was used for epithelial transport and metabolism studies, whereas the EpiIntestinal^TM^ Full tissue (FT; #SMI-200-FT) model was used for barrier, viability, and cytokine secretion experiments. After tissue delivery, the barrier integrity was assessed by transepithelial electrical resistance (TEER) measurements (paragraph 2.6) to confirm intact barrier function.

### 2.3. Cell viability

The impact of ENNs and BEA exposure on intestinal cell metabolic activity/viability was determined using the Alamar Blue assay according to the manufacturer’s protocol (Invitrogen, Thermo Fisher Scientific). The dark blue oxidised form of Alamar Blue (resazurin) is reduced to a highly fluorescent form (resorufin) by mitochondrial or cytoplasmic enzymes in living cells (O’Brien et al. 2000; Solhaug et al. 2025). The measured fluorescence intensity is thus proportional to the number of viable cells. Fluorescence (Excitation (Ex) 555 nm /Emission (Em) 585 nm) was quantified using the Spectramax i3x plate reader (Molecular Devices, San Jose, CA, USA).

### 2.4. Measurement of tissue thickness

After exposure, the EpiIntestinal^TM^ FT tissues were washed once in phosphate-buffered saline (PBS), fixed in 10% formalin for 48 h, embedded in paraffin blocks, and cut with a microtome (model HM3555, Thermo Scientific) at room temperature. Tissue sections were mounted on glass slides, and paraffin was removed by deparaffinization in 100% xylene, followed by rehydration through graded ethanol solutions (100%, 96%, and 70%) and rinsing in distilled water. The sections were subsequently stained with hematoxylin and eosin (H&E) according to the established protocol (Culling et al. 1985). H stains nuclei purplish blue, and E stains the extracellular matrix and cytoplasm pink. The slides were scanned using a microscope scanner (Nanozoomer S210, Hamamatsu Photonics, Hamamatsu, Japan) and analysed with NDP.view 3 software (*v*2.7.25; Hamamatsu Photonics). The tissue thickness was determined manually using 200 measurements per sample at 400 nm intervals.

### 2.5. Cytokine analysis

*ELISA:* After exposure, basolateral medium was harvested and analysed for secreted IL-8 and IL-6 by enzyme-linked immunosorbent assay (ELISA), according to the manufacturer’s guidelines (Human IL-8 CytoSet^TM^ #CHC1303 and Human IL-6 Matched Antibody Pair #CHC1263, respectively; Life Technologies, Invitrogen, Thermo Fisher Scientific). The absorbance at 450 nm was measured using a plate reader (Spectramax i3x). *Cytokine bead assay (CBA):* After exposure, basolaterally harvested cell culture medium was simultaneously measured for several secreted cytokines (IL-1β, TNF, IL-6, IL-8, IL-10, IL-12p70) using the BD Cytometric Bead Array (CBA), human inflammatory cytokine kit (#551811; BD Biosciences, Thermo Fisher Scientific) as described by the manufacturer, and analysed by flow cytometry (Ex 633 and 488, Em FL4 (675/25) vs FL2 (585/40)) using an Accuri C6 flow cytometer (BD Biosciences, Thermo Fisher Scientific).

### 2.6. Epithelial barrier integrity

Barrier integrity in the different models was assessed by measuring transepithelial electrical resistance (TEER) using a Millicell RES Voltmeter (Millipore, Burlington, MA, USA).

### 2.7. Secretion of mucin glycoprotein MUC5AC

Detection of secreted MUC5AC, a major gel-forming mucin produced by surface epithelial goblet cells, was performed using a protocol modified from Wan et al. (2014). Apical cell culture medium of Caco-2/HT29-MTX-E12 co-culture or EpiIntestinal^TM^ FT tissues was diluted 1:4 in cell culture medium and then mixed 1:2 with coating buffer (sodium carbonate 0.16 M/hydrogen carbonate 0.034 M; pH 9.6) in an ELISA plate (Maxisorb, Thermo Fisher Scientific) in a total volume of 100 µL, and dried at 40 °C overnight. The ELISA plate was then washed twice with PBS/Tween-20, 0.05% (PBST), followed by blocking with assay buffer (PBST containing 2% bovine serum albumin (BSA) (Sigma-Aldrich). After 1 h, the plate was washed three times with PBST and incubated for 2 h with MUC5AC antibody (#MA1-21907, Invitrogen, Thermo Fisher Scientific) diluted 1:1000 in assay buffer on a plate shaker (700 rpm). The plate was washed three times with PBST and subsequently incubated with horseradish peroxidase (HRP)-conjugated goat anti-mouse antibody (1:2000; #7076, Cell Signaling Technology, MA, USA). After 2 h on a plate shaker (700 rpm), the plate was washed three times with PBST, and the colour reaction was developed with 3,39,5,59-tetramethylbenzidine (TMB) peroxidase solution (Neogen, Lansing, MI, USA) and stopped with 10% H_2_SO_4_. The absorbance at 450 nm was measured using a plate reader (Spectramax i3x).

### 2.8 Epithelial transport

Differentiated Caco-2 cells or EpiIntestinal™ tissues (SMI-100) in standing cell culture transwell inserts, as described in section 2.2, were placed in 12-well plates with 300 µL cell culture medium in the apical compartment and 1500 µL in the basolateral compartment. The cells were exposed apically with 5 µM ENNs and BEA and incubated for 4 h, with or without the addition of 5 µM P-gp inhibitor PSC833 (Sigma-Aldrich). Aliquots (150 µL) of the apical and basolateral medium were precipitated by mixing (1:1; v/v) with ice-cold acetonitrile (ACN), centrifuged for 10 min at 20,000×*g* and 4 °C, and the supernatants stored at −20 °C until analysis. For intracellular toxin analysis, the cells in each transwell insert were lysed with 100 µL CellLytic™ (Sigma-Aldrich) and homogenised by centrifugation through QIAshredder biopolymer shredding spin columns (Qiagen, Hilden, Germany) for 2 min at 4 °C and 20,000×*g* before protein precipitation with ACN as described above. The toxin distribution between the apical (AP) and basolateral (BL) compartments as well as the intracellular concentrations were determined by ultra-high-performance liquid chromatography high-resolution tandem mass spectrometry (UPLC–HRMS/MS) as described in section 2.10.

### 2.9 ENNB and ENNB1 biotransformation in EpiIntestinal™ tissues

EpiIntestinal™ tissues (SMI-100) in standing transwell inserts were placed in 12-well plates and apically exposed to 5 µM ENNB or ENNB1 for 4 h and prepared for UPLC-HRMS/MS as described in paragraph 2.8. Before analysis, two independent incubations were combined and up-concentrated to allow detection of minor metabolites. More specifically, the two replicates taken from the AP and BL compartments, as well as the cellular fraction, were pooled (each 150 µL, complemented with water if necessary). Subsequently, the samples were extracted by liquid–liquid extraction (LLE), adding 500 µL ethyl acetate and vortexing for 10 sec. The extraction was repeated three times, and the combined organic phases were evaporated to dryness under a gentle nitrogen stream at 40 °C. The residues were redissolved in 50 µL 50% ACN in water and stored at −20 °C until analysis. For verification, metabolite mixtures generated in human liver microsomes (HLM) on two separate days were included in the study and analysed for known ENNB (Ivanova et al. 2011) and ENNB1 (Ivanova et al. 2019) metabolites by UPLC–HRMS/MS.

### 2.10 Quantitative UPLC–HRMS/MS analysis of ENNs and BEA

Calibration curves were constructed by diluting mycotoxin stock solution in 50% ACN to the intended concentrations. Matrix-assisted calibration samples were produced by diluting culture medium (“matrix”) of Caco-2 cells or EpiIntestinal^TM^ tissues with ACN (1:1). The matrix preparations were stored for 20 min at −20 °C and centrifuged at 20,000×*g* at 4 °C for 1 min through 0.22 µm nylon Costar Spin-X filters (Corning Inc., Corning, NY, USA). After filtering through 0.45 µm Thomson Single StEP nano filters (Restek, Bellefonte, PA, USA), aliquots (40 µL) were transferred into HPLC vials and dried under a gentle stream of nitrogen at 50 °C. Subsequently, the dried matrix aliquots were reconstituted with 40 µL of solvent calibration samples to generate matrix-assisted calibration curves. On the day of analysis, all samples were thawed on ice and centrifuged again if precipitation was observed. The supernatants were filtered through the 0.45 µm Thomson Single StEP nano filters, and the AP samples were additionally diluted tenfold with 50% ACN, before all the samples were transferred into HPLC vials (40 µL aliquots) and placed into the UPLC autosampler. The UPLC–HRMS/MS analyses were performed using a Q-Exactive™ Hybrid Quadrupole-Orbitrap mass spectrometer equipped with a heated electrospray ion source (HESI-II), connected to a Vanquish Horizon ultrahigh-performance liquid chromatography (UPLC) system (ThermoFisher Scientific). The HESI-II interface was operated at 300°C in positive mode, with the following parameters: spray voltage 3.2 kV, capillary temperature 280 °C, sheath gas flow rate 35 l/min, auxiliary gas flow rate 10 l/min, and S-lens RF level 55. Analyses were performed in full-scan (FS) mode, including targeted fragmentation (data-dependent MS/MS, dd-MS^2^) of ions specified in an inclusion list. FS settings were as follows: full width at half maximum (FWHM) resolution, 70,000; automatic gain control (AGC) target, 1e6; maximum injection time (IT), 200 ms; scan range, *m/z* 600–950. The setting for the dd-MS^2^ was optimised as follows: FWHM resolution, 17,500; AGC target, 5e4; maximum IT, 150 ms; isolation window *m/z* 1.6; higher-energy collisional dissociation (HCD) with stepped normalised collision energy (NCE), 20, 35 and 60%. Chromatographic separation of the mycotoxins was achieved on a Phenomenex Kinetex^®^F5 column (2.1 × 100 mm, 1.7 µm) with 0.5 μm × 0.004 in. ID HPLC KrudKatcher Ultra Column In-Line filter (Phenomenex, Torrance, CA, USA) maintained at 30 °C. Water and ACN, both containing 2 mM ammonium formate and 2 mM formic acid, were used as mobile phases A and B, respectively. The flow rate was 250 μL/min and the injection volume 1 μL. The mycotoxins were eluted with a gradient starting at 5% B, increasing to 40% B in 5 min, remaining at 40% B until 7 min, then increasing to 70% B at 19 min, and finally increasing to 100% B at 23 min. The column was kept at 100 % B for 4 min before it was returned to the start conditions and re-equilibrated for 3 min. The total run time was 30 min. The quality of the quantitative UPLC–HRMS/MS method for ENNs and BEA was characterised by determining key performance parameters including the sensitivity, selectivity, linearity, accuracy and precision. Calibration curves were prepared using mycotoxin standards in 50% ACN and in cell culture medium (“matrix”). The method sensitivity was defined by the limits of detection (LOD) and quantification (LOQ), calculated as LOD = 3.3SD/s and LOQ = 10SD/s, based on the standard deviation of the y-intercept and the slope of the average regression line of two calibration sets analysed on the same day (Shrivastava and Gupta 2011). The obtained LOD and LOQ showed that the method was sensitive enough for the purpose of this study, ranging from 8 nM for ENNB in solvent to 154 nM for BEA in solvent (**Table 1)**. Selectivity was established by assessing the signal suppression/enhancement (SSE) of the matrix on the measured results. The SSE was determined for each mycotoxin by comparing the means of matrix-assisted and solvent standard calibration curves. The ratio of the curve slopes, expressed in percent, represented the SSE (SSE% = slope of matrix-assisted calibration curve/slope of solvent calibration curve*100 %), with values below 100 % indicating signal suppression and above 100 % enhancement. The results showed that the matrix had little effect on the UPLC-HRMS/MS analysis, except for BEA (**Table 1)**. Study samples were therefore quantified by using calibrations in solvent.

**Table 1.** Performance parameters of the multi-mycotoxin UPLC-HRMS/MS method including limits of detection (LOD, ng/mL) and quantification (LOQ; ng/mL) in solvent and in matrix-assisted calibration, and the relative signal suppression/enhancement (SSE) by the matrix.

| **Compound** | **Molecular formula** | **[M+NH<sub>4</sub>]<sup>+</sup> (m/z)** | **RT min** | **LOD\* (ng/mL)** |  | **LOQ\* (ng/mL)** |  | **SSE%\*** |
|---|---|---|---|---|---|---|---|---|
|  |  |  |  | **Solvent** | **"Matrix"** | **Solvent** | **"Matrix"** |  |
| ENNB | C<sub>33</sub>H<sub>57</sub>N<sub>3</sub>O<sub>9</sub> | 657.4433 | 16.1 | 5.1/16.5 | 18.9/6.8 | 15.6/49.9 | 57.2/20.5 | 113/98 |
| ENNB1 | C<sub>34</sub>H<sub>59</sub>N<sub>3</sub>O<sub>9</sub> | 671.4590 | 17.0 | 5.0/16.6 | 17.7/2.1 | 15.3/50.4 | 53.7/6.4 | 117/100 |
| ENNA1 | C<sub>35</sub>H<sub>61</sub>N<sub>3</sub>O<sub>9</sub> | 685.4746 | 17.8 | 11.2/100 | 17.1/7.7 | 33.8/30.3 | 51.8/23.4 | 123/76 |
| BEA | C<sub>45</sub>H<sub>57</sub>N<sub>3</sub>O<sub>9</sub> | 801.4433 | 18.4 | 40.1/22.3 | 19.9/9.3 | 121/67.7 | 60.4/28.3 | 172/105 |
| ENNA | C<sub>36</sub>H<sub>63</sub>N<sub>3</sub>O<sub>9</sub> | 699.4903 | 18.6 | 21.9/11.8 | 16.2/7.0 | 66.2/35.8 | 49.1/21.2 | 128/103 |

\*Caco-2 cells/EpiIntestinal<sup>™</sup> tissues

The linearity of the calibration curves (weighed 1/x) was evaluated using the regression coefficients R^2^, which were > 0.99 for all mycotoxins in both solvent and matrix (data not shown). Method accuracy was determined by calculating the recoveries of mycotoxin standards spiked into Caco-2 medium and comparing the nominal and measured levels in the range of 1 ng/mL to 500 ng/mL. Above the respective LODs, the recoveries of the four ENNs were within 100 ± 5% (n = 6 analyses), whereas for BEA a recovery of 100 ± 15% (n = 2) was achieved. The intermediate precision of the method, defined as relative standard deviation (%RSD), was below 10% for ENNs, and below 12% for BEA (**Supplementary Table 1**). For EpiIntestinal^TM^ medium, only two analyses could be performed due to limited material access. Recoveries for concentrations above the LOD were 100 ± 20% for ENNs and BEA. The results were highly reproducible with %RSD below 10% for all mycotoxins.

### 2.11 UPLC–HRMS/MS analysis of ENNB and ENNB1 biotransformation products

ENNB and ENNB1-related metabolites were determined using an Orbitrap Fusion Tribrid mass spectrometer (Thermo Fisher Scientific) equipped with a heated electrospray ion source (HESI-II) and coupled to a Vanquish Horizon ultrahigh-performance liquid chromatography (UPLC) system (Thermo Fisher Scientific). The HESI-II interface was operated in positive ion mode at 300 °C; other parameters were adjusted as follows: spray voltage 3.8 kV, heated transfer capillary temperature 280 °C, sheath gas flow rate 35 L/min, auxiliary gas flow rate 10 L/min, S-lens RF level 60. Date acquisition in FS mode was conducted with a resolution of 240,000 FWHM at an *m/z* range of 600 to 1000. For MS^2^ experiments, precursors were directly identified in the MS^1^ scan by utilising a targeted mass difference of 4.9554 Da between sodiated molecular ions and ammonia molecular ions. Fragmentation was achieved by collision-induced dissociation (CID) or higher-energy C-trap dissociation (HCD)of the sodiated molecular ions, which yielded the most informative MS^2^ spectra, which were recorded at 30,000 resolution with an isolation window of *m/z* 1.4. The maximum injection time was set to 80 ms for both fragmentation techniques. HCD collision energy was normalised and set to 50%, while CID collision energy was set to a fixed 35 %. The UPLC system was run under the same conditions as described above for the quantitative method.

Major ENNB and ENNB1 biotransformation products detected in the AP, BL or cell compartments of EpiIntestinal^TM^ tissues grown in the transwell system were tentatively annotated based on the [M+NH₄]⁺ and [M+Na]⁺ adducts using an in-house library. Data were processed in Compound Discoverer, version 3.3 (Thermo Fisher Scientific) and characterised by matching retention times (RT) (tolerance 0.15 min), masses (tolerance 5 ppm), and fragmentation data to the entries in a previously established in-house MzVault 2.3 SP1 library generated from incubations in HLM (Ivanova et al. 2024). Due to the low metabolite levels in the samples and the resulting lack of suitable HRMS² data for sodium adducts, the confidence in metabolite identification was generally at 60% and thus relatively low. Considering the overall low abundance of the detected metabolites, a semi-quantitative analysis (calculated using Xcalibur, version 4.2.28.14; Thermo Fisher Scientific) was conducted based on the sum of the [M+NH₄]⁺ and [M+Na]⁺ adducts of each metabolite. The calculated levels were subsequently used to show the distribution of the detected metabolites across the different cellular compartments.

### 2.12 Interaction of ENNB with the hepatic biotransformation of E2

The effect of ENNB on the hepatic metabolism of the sex hormone E2, i.e. the formation of E2-related metabolites such as estrone (E1) and 2-hydroxyestradiol (2-OH-E2), was investigated *in vitro* using commercially available human liver microsomes (HLM; No. X008067, Lot IHG; Celsis *In Vitro* Technologies, Baltimore, MD, USA). E2, E1 and 2-OH-E2 were purchased from Sigma-Aldrich and incubations were performed as previously described (Mallia et al. 2020). Briefly, E2 (5 µM) was incubated alone or with ENNB (0.5, 2, or 5 µM) in 0.5 mL 45 mM HEPES buffer (pH 7.4) containing 2 mg/mL HLM protein and an NADPH-generating system (0.91 mM NADPH, 0.83 mM NADP^+^, 19.4 mM glucose-6-phosphate, 1 U/mL glucose-6-phosphate dehydrogenase, and 9 mM MgCl_2_·6H_2_O) in a shaking water bath at 37 °C. Aliquots were withdrawn after 0, 5, 10, 15, 30 and 60 min, and the enzymatic reactions were stopped with the addition of ice-cold acetonitrile (1:1, v/v). The samples were centrifuged for 10 min at 20,000×*g* and 4 °C to precipitate proteins, filtered with 2,000×*g* for 1 min at room temperature through 0.22 µm Costar Spin-X filters, transferred to HPLC vials and analysed by quantitative UPLC-triple-quadrupole mass spectrometry (UPLC-3Q-MS/MS).

### 2.13 UPLC-3Q-MS/MS of E2 and its biotransformation products

E2 and E2-related compounds were determined using an Agilent 1290 Infinity Binary UPLC system coupled via an electrospray ionisation (ESI) interface to an Agilent 6470 TripleQ mass spectrometer (Agilent Technologies, Santa Clara, CA, USA). The targeted method was performed as previously described (Mallia et al. 2020), detecting definite precursor ion-to-product ion transitions for E2 (*m/z* 255.2 to *m/z* 159.0; RT 13.5 min), E1 (*m/z* 271.2 to *m/z* 133.0; RT 15.1 min), 2-OH-E2 (*m/z* 271.2 to *m/z* 175.0; RT 11.8 min) and 4-OH-E2 (*m/z* 271.2 to *m/z* 175.0; RT 12.0 min). Briefly, the ESI interface was operated in positive ion mode with the following MS parameters: drying gas temperature 280 °C, drying gas flow 10 L/min, nebuliser gas pressure 25 psi, sheath gas temperature 375 °C, sheath gas flow 12 L/min, nozzle voltage 1000 V and capillary voltage 4 kV. Compounds were separated using a 150 × 2.1 mm i.d. Kinetex F5 (2.6 µm) HPLC column, maintained at 30 °C, with a 0.5 µm KrudKatcher Ultra pre-column filter (Phenomenex) at a flow rate of 0.25 mL/min. Compounds (injection volume 1 µL) were eluted using water and ACN as mobile phases A and B, respectively, applying a gradient starting at 2 % B (0–6 min), increasing to 38 % B at 8 min and reaching 48 % B at 18 min. The total run time was 24 min, including a 3 min column washing step at 95 % B, followed by a 2 min re-equilibration step before the next injection. The quantitative analysis was performed using matrix-assisted calibrations, and data were processed with MassHunter (version 10.2; Agilent Technologies).

### 2.14. Statistical analyses

Experimental data were analysed using GraphPad Prism (version 9.0.1.151; GraphPad Software, Boston, MA, USA). Applying One- or Two-Way ANOVA followed by Dunnett’s post-hoc test, study results were considered as statistically significant at p < 0.05. In general, the number of independent experiments was greater than three. However, the high cost of EpiIntestinal^TM^ tissues limited their availability, and therefore, two independent incubations were performed for this cell model.

## 3. Results and Discussion

Concerns about potential low-level chronic toxicity of ENNs and BEA is based on the high prevalence of these mycotoxins in cereal-based food products, and on the wide range of effects reported in numerous *in vitro* experiments. ENNs and BEA have been attributed with antibacterial, antifungal, antiviral, anticancer, antiparasitic and insecticidal properties, affecting oxidative stress, apoptosis, inflammation, cholesterol synthesis, platelet activation, nervous and endocrine systems (Caloni et al. 2020; Feudjio et al. 2010). The ionophoric nature of these mycotoxins enables cation transport across cell membranes, which is closely linked to their broad range of cellular targets. There have been made attempts to exploit the pharmacological potential of ENNs and BEA; however, the *in vivo* efficacy has been found deficient. Thus, a considerable gap exists between the known consequences of exposure in cellular test systems and those observed in humans and animals after oral exposure. Factors contributing to this discrepancy include the intra-intestinal bioaccessibility and toxicokinetic characteristics of the mycotoxins, considering that low absorption, high distribution, biotransformation and excretion can limit the level of active molecules at the target sites (Manzini et al. 2015; Meca et al. 2012). Moreover, the most relevant mechanisms of action for chronic toxicity remain undefined. In this context, the present study has addressed important aspects regarding this knowledge gap by determining the effects of ENNs and BEA in the gut, the first point of contact of the mycotoxins with epithelial surfaces after release from the food matrix. Particularly, their impact on the gastrointestinal epithelium, intestinal immune system and biotransformation were examined in human intestinal cells, as well as potential interference with the metabolism of endogenous E2, possibly leading to disturbances of hormone homeostasis.

### 3.1 Predicted intra-intestinal ENNs and BEA exposure after dietary uptake and relevance of cell studies

Data compilations in reviews evaluating the presence of ENNs in European cereals and derived foods show that median concentrations usually range from 3.2 to 166 µg/kg, but that in individual batches levels of up to 635 mg/kg can be reached (Coulet et al. 2025; Gautier et al. 2020). Based on measured ENNs and BEA concentrations in European cereal-based foods and consumption data of different age groups, EFSA estimated the mean dietary intake of ENNs (sum of ENNB, ENNB1, ENNA, and ENNA1) to 0.42 – 1.82 µg/kg body weight (bw)/day (EFSA 2014). High intake (95^th^ percentile) was estimated to 0.91 – 3.28 µg/kg bw/day, and worst-case acute intake as high as 4.67–10.1 µg/kg bw/day (EFSA 2014). For BEA, the mean dietary intake was calculated to 0.003-0.050 µg/kg bw/day, the high intake to 0.006– 0.093 µg/kg bw/day, and the acute intake to 0.05–0.1 µg/kg bw/day. Notably, toddlers had the highest estimated intake of both ENNs and BEA because they eat more in proportion to their body weights (EFSA 2014). From these intake estimates, possible concentrations of ENNs and BEA in the human gastrointestinal tract of small children after a meal were estimated: Assuming a body weight of 20 kg and a stomach volume of 0.5 mL for toddlers, distributing the daily intake equally over 3 meals and using the highest estimates of mean and high intakes, the concentrations in the stomach after a meal would amount to 0.2 µM ENNs and 0.002 µM BEA. These levels could also increase by about 10-fold in the small intestine due to water reabsorption, but also decrease due to limited bioaccessibility and in-gut fermentation (Pallarés et al. 2020). Despite these uncertainties, the prediction indicated that the ENNs concentrations in the low micromolar range used in this study were realistic. In contrast, the actual exposure to BEA is apparently well below the concentrations used in this study.

### 3.2 ENNB-induced responses in Caco-2/HT29-MTX-E12 co-culture

The molecular ring structure of ENNs and BEA enables them to form transmembrane, cation-selective channels, conferring ionophoric properties that affect organelles such as lysosomes and mitochondria, the plasma membrane, and cation-dependent physiological pathways (EFSA 2014). The ability to form ionophores is thus considered the primary mode of action (Behr et al. 2025; Dayeh et al. 2025), and the major cause of observable toxicological outcomes such as the destabilisation of lysosomes (Gammelsrud et al. 2012; Ivanova et al. 2012; Oliveira et al. 2020).

ENNB, the so far most-studied ENN (Behr et al., 2025), was selected for preparatory experiments in this study, which aimed at optimising culture conditions, identifying non-cytotoxic concentrations and exploring the impact on important endpoints such as general toxicity, barrier integrity, cytokine secretion and mucus production. In this study, experiments were performed with a differentiated co-culture of Caco-2/HT29-MTX-E12 cells grown on transwell inserts. After about 21 days, the co-culture is differentiated into a polarised epithelial monolayer with tight junctions, forming a functional barrier similar to the intestinal lining, including brush border microvilli, digestive enzymes and multiple transporters for nutrient uptake and drug transport (Behrens et al. 2001; Béduneau et al. 2014; Pan et al. 2015).

Cell viability, determined by metabolic activity using the Alamar Blue assay, was not significantly affected by ENNB exposure at concentrations up to 10 µM for 24 h (**Fig. 2A**). Information on ENNB-induced toxicity in fully differentiated monolayers is still very limited, but it has been reported that differentiated Caco-2 cells are more resistant to external stressors than dividing cells (Bu et al. 2016). This also aligns with previous studies using undifferentiated Caco-2 cells, where ENNB in the range of 2.5–30 µM, reduced cell viability after 24 h by 20 % (Fernandez-Blanco et al. 2016; Meca et al. 2011) or 50% (Ivanova et al. 2012), measured by metabolic activity assay (MTT) or lysosome integrity assay (Neutral Red), respectively. Furthermore, in partly differentiated Caco-2 cells, ENNB (5 µM) combined with IL-1β (25 ng/mL) reduced metabolic activity by about 30% after 20 h of exposure, as measured by the CellTiterBlue assay (Elesh et al. 2026). In contrast to fully differentiated Caco-2 monolayers that resemble mature enterocytes in both structure and function, proliferating and partially differentiated Caco-2 cells may represent proliferative intestinal crypt cells that have a higher sensitivity to ENNB. Both models are highly relevant but represent different parts of the intestine.

**Fig. 2.**
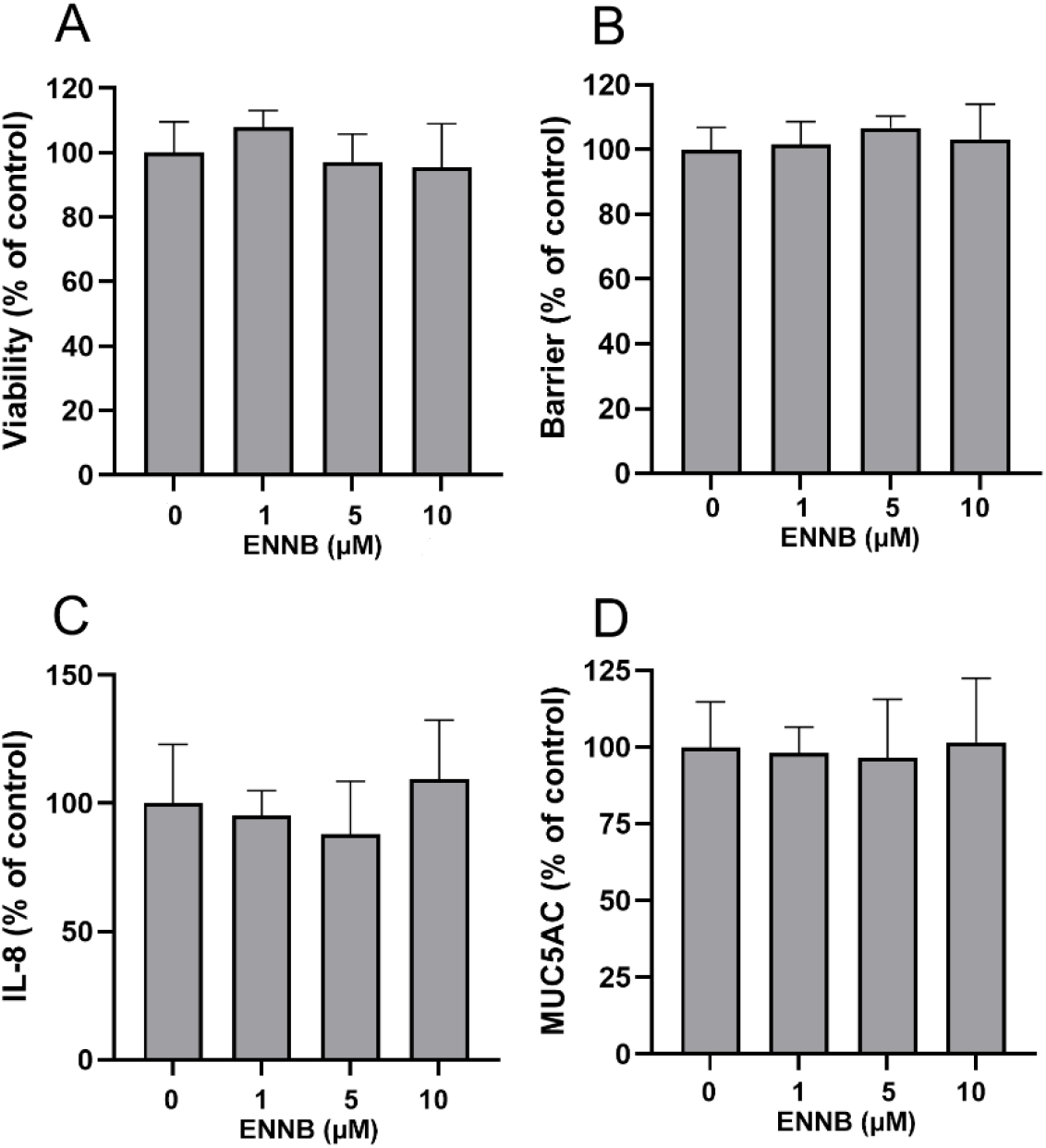
Effects of ENNB in Caco-2/TH29-MTX-E12 co-cultures treated for 24 h; (**A**) viability (metabolic activity), (**B**) barrier integrity (TEER), (**C**) IL-8 secretion and (**D**) MUC5AC secretion. Data are expressed as mean ± SD of 5 independent experiments. Significant differences (p< 0.05) as compared to ctrl are indicated with an asterisk (*). One-way ANOVA with Dunnett’s post hoc test.

Barrier integrity of the Caco-2/HT29-MTX-E12 co-culture grown on transwell inserts was evaluated by measuring the TEER, which reflects the resistance to ion passage across monolayers. When the co-culture is fully differentiated, the TEER values reach a plateau, normally observed after 18 to 21 days (Pan et al. 2015). Potential disruption of the barrier functionality caused by ENNB exposure, visible through reduced TEER values because of a more permeable cell layer, was measured at concentrations up to 10 µM for 24 h, showing no notable effect (**Fig. 2B**). These findings also supported that the differentiated monolayer was more resistant to ENNB exposure than what had been previously observed in dividing cells (Behr et al. 2025). Fittingly, a recent study in differentiated Caco-2 cells showed that exposure to ENNB1 at concentrations up to 6 µM for 24 and 48 h did not decrease the TEER, although the mycotoxin induced some changes in the intercellular junctional organisation (Ranaldi et al. 2026).

Intestinal epithelial cells are part of the mucosal immune defence and produce cytokines and chemokines that recruit and activate immune cells (Pitman and Blumberg 2000). Enterocytes secrete IL-8 to attract neutrophils in response to inflammatory stimuli, particularly during acute inflammation, but also under chronic inflammatory conditions (Eckmann et al. 1993; Herrero-Cervera et al. 2022; Sonnier et al. 2010). Elevated IL-8 levels have also been observed in inflammatory bowel disease (IBD), where they contribute to sustain intestinal inflammation (Izutani et al. 1995). Interestingly, the IL-8 secretion in the Caco-2/TH29-MTX-E12 co-culture remained unaffected by ENNB exposure for 24 h at concentrations up to 10 µM (**Fig. 2C**). In contrast, a recent study exposing differentiated Caco-2 cells to ENNB1 (0.75-6 µM, 24 h and 48 h) showed increased IL-8 secretion already at the lowest concentration (Ranaldi et al. 2026). Although no dose-response relationship was observed, ENNB1 appeared to induce a proinflammatory response in the cells despite no measurable effects on barrier integrity. Furthermore, a recent study using partly differentiated Caco-2 cells showed that incubation with 1 µM ENNA or ENNB1 in the presence of IL-1β (25 ng/mL) induced a slight increase in cytokine gene expression compared with IL-1β alone (Elesh et al. 2026). Specifically, ENNA increased IL-6 and IL-10 mRNA levels, whereas ENNB1 increased IL-6 and IL-8. In contrast, treatment with ENNB, ENNA1, or BEA showed no significant changes. Nevertheless, inflammatory endpoints remain rather understudied for ENNs and BEA, highlighting the need for further investigations.

The mucus lining the intestinal epithelium is an important physicochemical barrier that provides protection against ingested pathogens and toxins (Grondin et al. 2020). Mucins are essential components of the mucus layer, regulating viscosity and contributing to antimicrobial defence (Sheng and Hasnain 2022). Significant changes in mucin levels have been associated with gastrointestinal diseases like IBD (Kang et al. 2022). Although MUC2 is the dominant secreted gel-forming mucin in the intestinal mucus layer, increased secretion of the gel-forming MUC5AC is a typical marker for inflammatory or injury-related conditions (Olli et al. 2020). Since the TH29-MTX-E12 cells produce mucus, the co-culture is suitable for assessing mycotoxin-induced effects on mucin expression. Thus, several studies have shown that mycotoxins can affect the mucus layer (Robert et al. 2017), such as the *Fusarium* mycotoxin deoxynivalenol, which was found to upregulate the mRNA expression of several mucins in Caco-2 cells (Wan et al. 2025), and increase MUC5AC and MUC5B secretion in the Caco-2/HT29-MTX co-culture (Wan et al. 2014). However, in the present study ENNB concentrations up to 10 µM did not have any impact on the secretion of MUC5AC in the co-culture monolayer after 24 h exposure (**Fig. 2D)**.

### 3.3 Effects of ENNs and BEA in human primary intestinal cells, the EpiIntestinal tissues

Based on findings from the Caco-2/HT29-MTX-E12 co-culture (section 3.2) and estimates of real-life concentrations in the gastrointestinal tract (section 3.1), concentrations of up to 10 µM were used in experiments with human primary EpiIntestinal™ tissues. Concentration-dependent effects were assessed in two independent replicates for all mycotoxins. Since the number of mucus-producing cells in EpiIntestinal ^TM^ FT tissue is rather limited (Personal communication, 2024; Markus J, Mattek Life Sciences), mucus secretion was not measured in this cell system.

#### 3.3.1 Effects on viability and barrier

ENNB (1–10 µM) and ENNB1 (10 µM) significantly increased the metabolic activity/viability in EpiIntestinal^TM^ FT tissues after 24 h of exposure (**Fig. 3A**). Additionally, BEA slightly increased the metabolic activity/viability at 1 µM, but not at higher concentrations, while ENNA and ENNA1 did not have significant effects. This increase in metabolic activity, the opposite outcome of the decrease in cell viability usually seen at higher concentrations, could be evidence for the immediate cellular stress response to the presence of low toxin levels, indicating the activation of defence mechanisms (Solhaug et al. 2025; Tan et al. 2018).

**Fig. 3:**
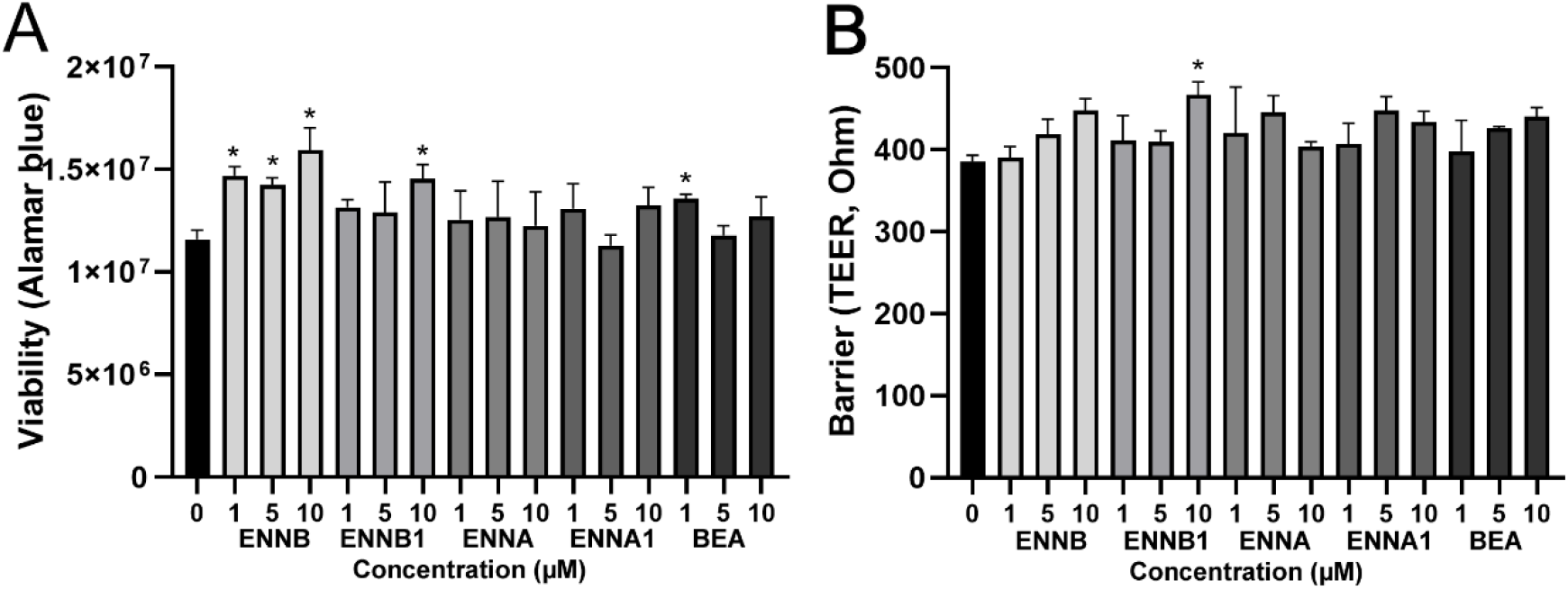
Effects of ENNs and BEA exposure (24 h) on **A**) viability (metabolic activity) and **B**) barrier integrity in EpiIntestinal^TM^ FT tissues. Data are presented as the mean ± SD of 2 independent incubations. Significant differences (p< 0.05) compared to ctrl are indicated with an asterisk (*). One-way ANOVA with Dunnett’s post hoc test.

Consistent with the results in the Caco-2/HT29-MTX-E12 co-culture (section 3.2), neither the ENNs nor BEA caused barrier rupture (TEER decrease) in the EpiIntestinal^TM^ FT tissues (**Fig. 3B**). Instead, a small increase in TEER at the highest ENNB1 concentration (10 µM) was observed, which would be consistent with an increased transcription of the tight junction protein occludin, as reported in rats exposed to single oral dose of an ENNs mixture (Cimbalo et al. 2021). In contrast, feeding pigs with a diet containing 2570 µg/kg BEA and 1345 µg ENNB + ENNB1/kg for 7 or 14 days did not affect zonulin transcription (Novak et al. 2021). However, the absence of an effect on zonulin does not exclude alteration in occludin, as zonulin acts upstream of proteins involved in intestinal barrier function (Sturgeon and Fasano 2016). However, all mycotoxin-containing diets led to moderate to severe histological lesions in the pig jejunum, so the applied feed concentrations might already have been high enough to cause acute toxic effects that would have made observing regulatory mechanisms difficult. Interestingly, ENNB1 (6 µM) affected intercellular junctional organisation in differentiated Caco-2 cells, particularly weakening adherens junctions, but without causing detectable changes in the TEER (Ranaldi et al. 2026). However, because lipophilic ENNs are more likely to be absorbed transcellularly than paracellularly, changes in junction proteins may not affect the absorption efficiency of these toxins.

Histological staining of ENNB-exposed EpiIntestinal^TM^ FT tissues showed a minor, but significant increase in tissue thickness at 5 µM, and a reduction at higher concentrations (20 and 30 µM) (**Fig. 4**). The slight tissue thickening at the non-toxic, lower level could have been caused by stress-induced cellular swelling associated with reversible epithelial injury, which is consistent with the observed increase in metabolic activity (**Fig. 3A**). The reduction in tissue thickness at higher ENNB levels could, however, reflect toxicity and loss of cellular integrity, aligning well with the reported tissue damages in the jejunum of pigs that had received a diet containing substantial amounts of ENNB and ENNB1 (Sturgeon and Fasano 2016).

**Fig. 4:**
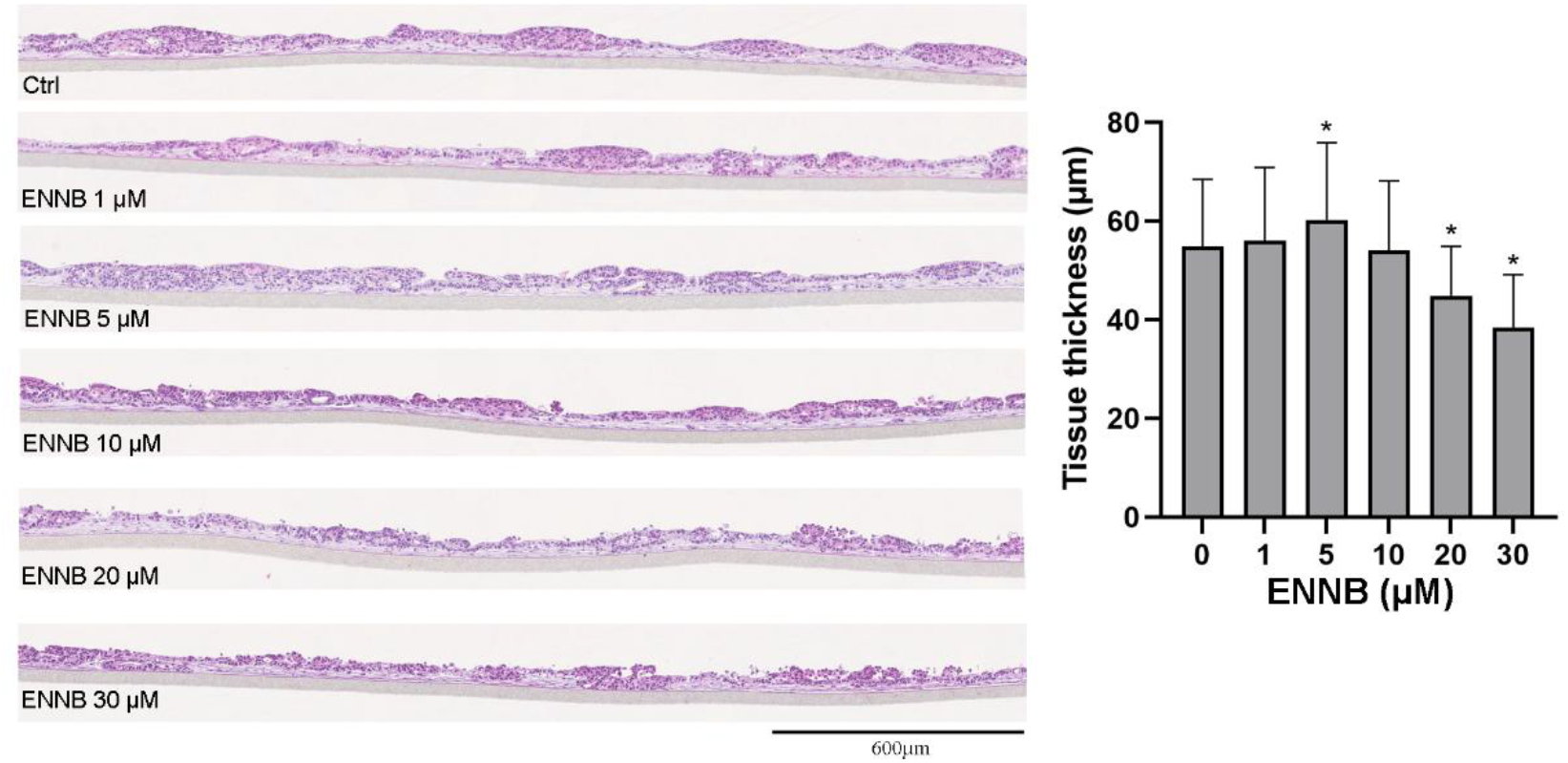
Effects of ENNB (24 h) exposure on the thickness of EpiIntestinal^TM^ FT tissues. Data are presented as mean ± SD from 2 independent incubations, with approximately 200 measurements of the tissue thickness at 400 nm intervals for each exposure condition. Significant differences (p< 0.05) compared to ctrl are indicated with an asterisk (*). One-way ANOVA with Dunnett’s post hoc test.

#### 3.2.2 Secretion of cytokines

To assess potential inflammatory responses in EpiIntestinalFT tissues, IL-1β, TNF, IL-6, IL-8, IL-10, and IL-12p70 were initially screened (**Supplementary Fig. 1**). This panel was selected to cover different aspects of intestinal inflammatory signalling. *In vivo*, intestinal epithelial cells respond to pathogenic bacteria and cytokine signals from neighbouring immune and stromal cells, including fibroblasts. The EpiIntestinalFT model incorporates epithelial cells and an underlying fibroblast-containing compartment, thereby allowing epithelial-stromal interactions to contribute to the inflammatory response. Increased secretion of IL-6 and IL-8 indicated a proinflammatory effect of ENNs and BEA, even though the other cytokines (IL-1β, TNF, IL-10, and IL-12p70) included in the multi-method showed no visible change. However, these cytokines are predominantly produced by activated innate immune cells, including monocytes, macrophages and dendritic cells (Abbas et al. 2025), which are not present in the EpiIntestinal™ FT model. Based on the initial screening, IL-6 and IL-8 were selected for further analysis, representing a pleiotropic inflammatory cytokine and a chemokine involved in neutrophil recruitment, respectively.

In the next step, IL-6 and IL-8 secretion were specifically measured. As already seen in the Caco-2/HT29-MTX-E12 (**Fig. 2C**), ENNB did not increase the IL-8 secretion in the EpiIntestinal™ FT model (**Fig. 5A**). Comparable findings have been reported in other cell systems such as a co-culture consisting of bovine mammary epithelial cells (MAC-T) and macrophages (BoMac), where 29.9 µM ENNB, equivalent to the level eliciting 20% cytotoxicity (IC_20_) in the model, did not affect the IL-8 secretion (Shandilya et al. 2023). The slight IL-8 decrease in EpiIntestinal™ FT tissues at 1 µM may be linked to the observed increase in metabolic activity (**Fig. 3A**), which may be driven by activated cellular defence mechanisms. Similarly, only a modest, but non-significant reduction in IL-8 mRNA expression was observed in undifferentiated intestinal epithelial cells (HCEC-1CT) following exposure to 0.1 or 1 µM ENNB in combination with IL-1β (25 ng/mL) (Elesh et al. 2026).

**Fig. 5.**
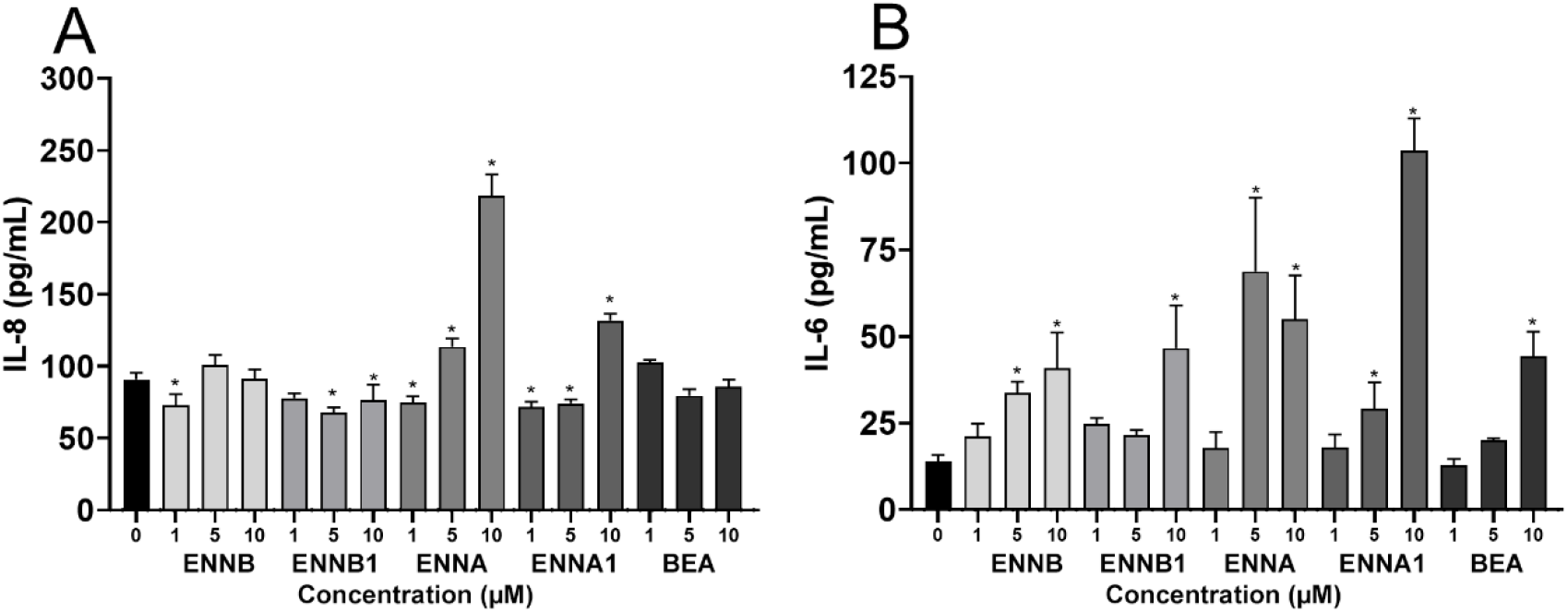
Effects of ENNs or BEA exposure (24 h) on cytokine secretion, IL-8 (**A**) and IL-6 (**B**) in EpiIntestinal^TM^ FT tissues. Data are presented as mean ± SD from 2 independent incubations, each measured in triplicate. Significant differences (p< 0.05) compared to ctrl (0 µM) are indicated by an asterisk (*). One-way ANOVA with Dunnett’s post hoc test.

Exposure of the EpiIntestinal™ FT tissues to 5 µM and 10 µM ENNB1 led to a moderate but significant decrease in IL-8 secretion, which resembled the situation observed for ENNB (**Fig. 5A**). Interestingly, a recent study reported that ENNB1 induced an approximately twofold increase in IL-8 secretion in the AP compartment of differentiated Caco-2 cells, whereas no changes were detected in the BL compartment (Ranaldi et al. 2026). However, basal IL-8 secretion was relatively high, and the increase did not show a clear dose-response relationship, which should be considered when evaluating the biological relevance of this finding. Furthermore, the Caco-2 cells were differentiated using a protocol different from ours, based on the asymmetric differentiation model described by Ferruzza et al. (2013). Thus, differences in basal cytokine secretion and differentiation conditions may explain the divergent findings and suggest that ENNB1’s effects on IL-8 secretion may be model-dependent. In undifferentiated HCEC-1CT and partly differentiated Caco-2 cells exposed to 1 µM ENNB1 for 2 h, followed by co-incubation with IL-1β (25 ng/mL) for an additional 3 h, no significant changes in IL-8 mRNA expression were observed in HCEC-1CT cells, whereas both cytokines were significantly upregulated in Caco-2 cells (Elesh et al. 2026). This contrasting response suggests that cell type and differentiation status may influence the immunomodulatory effects of ENNB1 under inflammatory conditions. Although these results cannot be directly compared with ours, as ENNB1 was not examined in the absence of IL-1β stimulation, they provide additional evidence for the immunomodulatory potential of ENNB1. Moreover, ENNB1 (0.5 to 3 µM) incubation of porcine alveolar macrophages for 24 h resulted in 2.7-to 5.3-fold increase of IL-8 mRNA expression, while at 6 µM, the increase was reduced to 3.6-fold, which probably reflected the onset of cytotoxicity (Frazzini et al. 2026).

Interestingly, contrast to ENNB and ENNB1, ENNA and ENNA1 showed a substantial increase in IL-8 secretion in the EpiIntestinal™ FT tissues (**Fig. 5A**). ENNA produced a greater effect than ENNA1. The same pattern had also been described in the IL-1β-stimulated Caco-2 cells but was much less defined in the HCEC-1CT, where the IL-8 mRNA expression was not significantly changed by ENNA1 (Elesh et al. 2026). The different potencies of the four analysed ENNs to influence IL-8 secretion are apparently connected to the specific functional groups in the otherwise identical molecular structures (**Fig. 1**). These are determinants of the specific lipophilicities of the respective molecules that in turn appear to have a great impact on elicited biological effects. This is evident when comparing the order of lipophilicity (i.e., ENNA>ENNA1>ENNB1>ENNB; (Tran et al. 2020) with the effect amount of IL-8 secretion in EpiIntestinal™ FT tissues (this study) and Caco-2 cells (Elesh et al. 2026), which follow the same order.

BEA at levels up to 10 µM had apparently no effect on the IL-8 secretion in EpiIntestinal™ FT tissues (**Fig. 5A**), and similar outcomes, but at the level on mRNA expression, were found in Caco-2 cells exposed to 0.1 and 1 µM BEA in combination with IL-1β (25 ng/mL) (Elesh et al. 2026). This ineffectiveness may differ in epithelial cells from organs other than the intestine. E.g., in the bovine MAC-T/BoMac co-culture, exposure with 11.3 µM BEA (IC_20_) for 48 h decreased IL-8 significantly, with and without pre-treatment with immune defence-stimulating bacterial lipopolysaccharide (LPS). BEA is even more lipophilic than the ENNs (Tran et al. 2020); nevertheless, the reduced activity might be caused by its considerable structural difference, containing three benzyl rings in the molecule that change its spatial dimensions significantly and possibly affect the interaction potential with biological targets (**Fig. 1**).

IL-6 expression was induced by all ENNs and BEA in the EpiIntestinal™ FT tissues, although with different efficiencies (**Fig. 5B**). ENNB and ENNB1 exposure at 10 µM resulted in a 3-to 4-fold increase in IL-6 as compared to the untreated control, whereas ENNA and ENNA1 increased IL-6 about 5-to 9-fold, respectively. The effectiveness of 10 µM BEA was comparable to that of ENNB. For IL-8, the observed differences in potency again related to the compound ranking in lipophilicity, including BEA.

A similar pattern of ENN-induced changes in IL-6 was recently reported in IL-1β-stimulated, partly differentiated Caco-2 cells (Elesh et al. 2026). Exposure to 1 µM ENNB, ENNB1, ENNA, or ENNA1 increased IL-6 mRNA expression by 1.5-, 1.1-, 2.1-, and 1.3-fold, respectively. In contrast, in undifferentiated HCEC-1CT cells, the corresponding IL-6 mRNA levels were 0.5-, 0.4-, 1.1-, and 0.6-fold relative to control, respectively, indicating an overall reduction in IL-6 expression. Despite the difference in response direction, the relative potency of the ENNs was similar in the two cell models. Incubation with 1 µM BEA resulted in an IL-6 decrease in both cell models, and the same was observed for all mycotoxins at 0.1 µM. This clear dose-dependent effect was also observed in porcine alveolar macrophages exposed to ENNB1 for 24 h (Frazzini et al. 2026). Here, ENNB1 decreased IL-6 mRNA expression by 0.05-and 0.5-fold at 0.5 µM and 1.5 µM, respectively, while 3 µM ENNB resulted in a 13-fold increase. Furthermore, in the bovine MAC-T/BoMac cell culture, exposure with 11.3 µM BEA (IC_20_) produced a slight increase in IL-6 levels (Shandilya et al. 2023), illustrating again the importance of the cell type and culture conditions for the results obtained, which complicates the direct comparison of experiments.

Based on evidence from the present and referenced studies, there is convincing evidence that the cyclic hexadepsipeptic mycotoxins ENNS and BEA have immunomodulatory properties. At certain levels they could exacerbate an ongoing inflammation, but at relatively low concentrations, they might counteract the activation of the inflammasome. However, no changes in the secretion of IL-1β were reported from rats given a single oral dose of a mixture of ENNs (Cimbalo et al. 2021), but in that study, other cytokines were not included, indicating that analysis of a single cytokine may not provide a sufficient basis for the evaluation of effects on inflammation. In contrast to this, a study done using the murine macrophage cell line, RAW264.7, shows that ENNB (2.5 – 10 µM) in combination LPS (25 ng/mL) can activate the inflammasome and secretion of IL-1β, while LPS and ENNB alone had only a minor effect (Gammelsrud et al. 2012). Furthermore, when mice with induced colitis were dosed intraperitoneally for 7 days with 1, 2 or 4 mg/kg bw BEA, the colonic TNF-α, IL-1β, IL-12 and IFN-γ levels decreased. Moreover, BEA reduced lymphocyte infiltration into the mouse colon, leading to a dose-dependent alleviation of colitis symptoms such as diarrhoea and decreased body weight gain (Wu et al. 2013). This anti-inflammatory potential of BEA is proposed to be caused by inhibition of key enzymes in the NF-κB pathway (Wu et al. 2018). Genes identified as part of the “Inflammation mediated by chemokine and cytokine signalling pathway” by Kyoto Encyclopedia of Gene and Genome (KEGG) analysis were enriched by exposure to BEA or ENNB in the bovine MAC-T/BoMac co-culture (Shandilya et al. 2023). Finally, *in vivo* studies with intravenous application of 1, 3 and 5 mg/kg bw/day ENNB or ENNB1 in pregnant mice for 4 days showed a significant downregulation of CXCL1, IL-1β and IL-8 mRNA in foetal liver extracts at the 5 mg/kg bw dose, demonstrating the potential to modulate or even suppress embryonic innate immune functions (Huang et al. 2022; Huang et al. 2019).

### 3.3 Uptake and metabolism of ENNs and BEA in intestinal cells

A toxin’s efficacy at the target site depends not only on its toxicological potency but also on its toxicokinetic properties. After oral uptake, the toxin must be released from the food matrix, absorbed from the gut into the bloodstream, and distributed throughout the organism to reach the action site. The fraction absorbed critically determines a compound’s bioavailability and correlates with its lipophilicity. For highly lipophilic molecules, such as ENNs and BEA (Tran et al. 2020) lipid affinity drives interactions with cell membranes and adhesion to non-polar surfaces. Understanding the extent of absorption and transport through intestinal epithelial cells is therefore essential for interpreting the effects observed in different cell models.

#### 3.3.1 Uptake and transport

Uptake studies were performed for the ENNs and BEA in differentiated Caco-2 cells and primary EpiIntestinal^TM^ tissues. For these experiments, the Caco-2 cells were not grown in co-culture with the mucus-producing HT29-MTX-E12 cells because the mucus could interfere with the determination of the compounds’ specific transepithelial transport ratios. Since transport studies should be performed under linear conditions and at non-toxic concentrations, all experiments in both cell systems were performed by incubating 5 µM of the respective mycotoxin for 4 h, based on an exploratory test using ENNB1 and Caco-2 cells (**Supplementary Fig. 2**). Transport ratios across the epithelium were measured from the luminal side (AP) into the systemic circulation (BL). The toxin distribution across the transwell system (AP, BL, cells) was determined by considering the measured concentrations and volumes in each compartment and calculating the proportions of the total amount (**Table 2**). Lipophilic compounds such as the ENNs and BEA could adhere to plastic surfaces during the experiments. To limit this, the plates were saturated with the FCS-containing culture medium for 30 min at 37 °C before the incubations. Additionally, compound amounts in all compartments of the transwell system were summarised after 4 h and compared to the added amount, showing approximate recovery rates in the range of 80-114 %, depending on the cell model and the mycotoxin (**Supplementary Table 2**).

**Table 2.** Distribution of the ENNs and BEA between the AP and BL compartments, and in the cells, in the Caco-2 and EpiIntestinal^TM^ models. The relative distributions are expressed as mean ± SD of 3-4 independent experiments (Caco-2) or as mean ± SD of 2 independent incubations (EpiIntestinal^TM^ tissues). Significant differences (p< 0.05) after co-incubation with the inhibitor PSC833 are indicated by asterisk (*). Two-way ANOVA with Sidak post hoc test.

|  | Toxin distribution (%) |  |  |  |  |  |
|---|---|---|---|---|---|---|
|  | Caco-2 |  |  | EpiIntestinal<sup>TM</sup> tissue |  |  |
| Toxin ± PSC833 | AP | BL | Cells | AP | BL | Cells |
| ENNB | 56 ± 1.7 | 42 ± 1.7 | 2.1 ± 0.1 | 57 ± 1.0 | 32 ± 1.8 | 12 ± 0.9 |
| ENNB + PSC833 | 56 ± 2.5 | 42 ± 3.1 | 2.1 ± 0.7 | 45 ± 0.9<sup>*</sup> | 45 ± 2.1<sup>*</sup> | 9.2 ± 1.2<sup>*</sup> |
| ENNB1 | 64 ± 1.8 | 33 ± 1.4 | 2.4 ± 0.5 | 64 ± 1.3 | 25 ± 1.2 | 11 ± 0.1 |
| ENNB1 + PSC833 | 62 ± 1.2<sup>*</sup> | 36 ± 1.3<sup>*</sup> | 2.7 ± 0.3 | 46 ± 1.6<sup>*</sup> | 43 ± 1.6<sup>*</sup> | 11 ± 0.03 |
| ENNA | 74 ± 3.1 | 22 ± 2.8 | 3.6 ± 0.8 | 72 ± 7.2 | 8.1 ± 0.4 | 20 ± 6.8 |
| ENNA + PSC833 | 71 ± 2.6 | 24 ± 1.9 | 4.8 ± 1.0 | 63 ± 1.7 | 15 ± 0.4 | 22 ± 2.0 |
| ENNA1 | 67 ± 2.5 | 30 ± 2.3 | 3.1 ± 0.3 | 76 ± 0.7 | 9.1 ± 0.3 | 15 ± 1.0 |
| ENNA1 + PSC833 | 65 ± 1.3 | 31 ± 0.7 | 3.6 ± 1.0 | 61 ± 2.1<sup>*</sup> | 22 ± 0.2<sup>*</sup> | 17 ± 1.9 |
| BEA | 75 ± 2.8 | 13 ± 1.3 | 12 ± 2.1 | 83 ± 0.6 | 6.1 ± 0.2 | 10 ± 0.4 |
| BEA + PSC833 | 73 ± 2.4 | 15 ± 2.5 | 11 ± 1.0 | 64 ± 0.7<sup>*</sup> | 11 ± 2.2 | 25 ± 1.5<sup>*</sup> |

In the Caco-2 cells, the ratios of transmembrane-transported ENNs and BEA after 4 h of AP exposure ranged from 13 % BEA to 42 % ENNB, in accordance with the suggested lipophilic ranking: BEA>ENNA>ENNA1>ENNB1>ENNB (Tran et al. 2020). Accumulation of the ENNs in the cells was 2 to 3%, whereas it was 12% for BEA (**Table 2**). In general, the uptake efficiency of compounds from the gut follows the empirical “Rule of five” for the intestinal permeability of compounds (Ro5), which states that a molecule missing any of the required conditions, i.e. molecular weight (MW) < 500 Da, octanol-water partition coefficient (log P) <5, number of hydrogen bond acceptors (HBA) < 10 and number of hydrogen bond donors (HBD) < 5, will not be well absorbed (Lipinski et al. 1997). The ENNs and BEA violate the MW (m/z, **Table 1**) and HBA criteria, and their predicted log P are close to 5 or, as for BEA, above (Tran et al. 2020). Apparently, the results obtained here in the Caco-2 cell model for the five mycotoxins were congruent with the Ro5, and similar findings have been previously reported (Behr et al. 2025). The transport ratios determined in different uptake models, including Caco-2 cells, porcine buccal mucosa cells and a human skin model, showed some variability depending on the incubation concentrations and times, but overall they assigned a higher transmembrane transferability to ENNB and ENNB1 than to ENNA, ENNA1 and BEA (Meca et al. 2012; Prosperini et al. 2012; Taevernier et al. 2015; Taevernier et al. 2016). Moreover, *in vivo* studies in rats and pigs orally dosed with an ENNs mixture confirmed the *in vitro* results (Devreese et al. 2014; Escrivá et al. 2015), demonstrating the applicability of the Caco-2 system to predict transport rates also for these mycotoxins.

It has been suggested that the ENNs and BEA are absorbed mainly via passive transcellular diffusion (Meca et al. 2012), consistent with their lipophilic and ionophoric properties (Krug et al. 2018). However, growing evidence also suggests that active transport mediated by efflux transporters such as P-gp (multidrug resistance protein-1; MDR-1, ABCB1), MDR-2 (ABCB4), and breast cancer resistance protein (BCRP, ABCG2) contributes considerably to the absorption ratios (Dornetshuber et al. 2009; Ivanova et al. 2010). Interestingly, using the Caco-2 cells, the P-gp inhibitor PSC883 caused a significant increase in ENNB1 of about 9% in the BL compartment (**Table 2**), consistent with the previous study by Ivanova and co-workers (Ivanova et al. 2010). However, because no effects were observed for the other ENNs and BEA, this may indicate mycotoxin specificity for individual transport proteins (Dornetshuber et al. 2009; Hiraga et al. 2005) or inadequate expression of these transporters in Caco-2 cells.

Consistent with the Caco-2 cell results, EpiIntestinal™ tissues showed the highest transport ratios for ENNB and ENNB1, while ENNA, ENNA1 and BEA permeated poorly from AP to BL (**Table 2**). However, intracellular toxin levels were considerably higher than in Caco-2 cells, with the highest accumulation found for ENNA, followed by ENNA1, ENNB, ENNB1, and BEA. In this case, the order of ENNs differed notably from the lipophilicity ranking, which may suggest the involvement of active transport mechanisms. In contrast to what was observed in the Caco-2 cells, PSC833 clearly inhibited the BL-to-AP efflux of ENNs and BEA in the EpiIntestinal^TM^ tissues, resulting in higher levels in the BL compartment (**Table 2**). The effect was the greatest for ENNB and ENNB1. The differences between the two cell models may, at least in part, be attributed to differences in the tissue architecture and composition. Whereas the Caco-2 epithelial monolayer is with about 25-30 µm relatively thin (Tan et al. 2018), the EpiIntestinal™ tissues are approximately 55 µm thick (**Fig. 4**), providing a physiologically relevant barrier that expresses a comprehensive set of ABC efflux transporters and biotransformation enzymes (Tan et al. 2018). It is therefore possible that the dynamic range of transmembrane compound absorption and transport differs between the two intestinal epithelium models.

Interestingly, co-incubation with the P-gp inhibitor PSC833 also considerably lowered the total amount of ENNs and BEA in the EpiIntestinal™ tissues incubation, while no difference was found in Caco-2 cells (**Supplementary Table 2**). A possible explanation can be that inhibition of P-gp-mediated efflux of the ENNs and BEA increases the intracellular retention of the mycotoxins, thereby prolonging their availability for metabolism by CYP3A enzymes, which are highly expressed in the EpiIntestinal^TM^ tissues but not in Caco-2 cells (Chothe et al. 2025; Küblbeck et al. 2016). This is also consistent with the proposed interplay between P-gp and CYP3A, whereby efflux transport regulates intracellular substrate concentrations and, consequently, the extent of metabolic conversion (Kirn et al. 1999).

#### 3.3.2 Metabolite profile of ENNB and ENNB1

The ENNs and BEA are substrates for CYP enzymes and mainly metabolised by hydroxylation, carboxylation and demethylation reactions (Behr et al. 2025). Large numbers of major and minor metabolites have been identified *in vitro* in incubations with biotransformation-competent liver fractions such as microsomes and primary hepatocytes (Fæste et al. 2011; Ivanova et al. 2019; Yuan et al. 2022), and in plasma samples of exposed animals and humans (Devreese et al. 2014; Fraeyman et al. 2016; Rodríguez-Carrasco et al. 2018). In contrast, no data are available on ENNs and BEA metabolism in the intestinal epithelium, partly because the usual intestinal model, i.e., Caco-2, is unsuitable for biotransformation experiments since the cells do not express sufficient CYP enzymes (Küblbeck et al. 2016). Thus, it was interesting to explore in the present study whether the EpiIntestinal™ tissues had retained the activity of their endogenous metabolic enzymes (Chothe et al. 2025) and could be used to investigate the intestinal biotransformation of mycotoxins.

ENNB and ENNB1 were added apically to the EpiIntestinal™ tissues, and after 4 h of incubation, the AP and BL compartments and the cells were analysed for parent compounds and derived metabolites. A previously developed optimised LC-HRMS/MS-tribrid workflow for metabolite detection and characterisation was used, which applied a novel scan method combining full scan analysis with two different fragmentation mechanisms, i.e. collision-induced dissociation (CID) and high-energy C-trap dissociation (HCD) (Ivanova et al. 2024). In total 13 metabolites of both ENNs were annotated with sufficient confidence in the different cell fractions based on observed ion masses and retention times aligned to an in-house metabolite library (**Table 3**; **Supplementary Tables 3 and 4**).

**Table 3.** Major ENNB and ENNB1-related metabolites detected in EpiIntestinal™ tissues after incubation for 4 h.

| ID | ΔRT* | RT ratio** | RT | Change | Formula | Confidence |
|---|---|---|---|---|---|---|
| **ENNB** |  |  |  |  |  |  |
| M1 | -4.8 | 0.737 | 13.1 | -(CH<sub>2</sub>) +(O) | C<sub>32</sub>H<sub>55</sub>N<sub>3</sub>O<sub>10</sub> | Low |
| M2 | -4.5 | 0.756 | 13.5 | -(H<sub>2</sub>) +(O<sub>2</sub>) | C<sub>33</sub>H<sub>55</sub>N<sub>3</sub>O<sub>11</sub> | Low |
| M3 | -4.1 | 0.774 | 13.8 | -(H<sub>2</sub>) +(O<sub>2</sub>) | C<sub>33</sub>H<sub>55</sub>N<sub>3</sub>O<sub>11</sub> | Low |
| M4 | -3.9 | 0.782 | 13.9 | -(CH<sub>2</sub>) +(O) | C<sub>32</sub>H<sub>55</sub>N<sub>3</sub>O<sub>10</sub> | Low |
| M5 | -3.9 | 0.786 | 14.0 | -(H<sub>2</sub>) +(O<sub>2</sub>) | C<sub>33</sub>H<sub>55</sub>N<sub>3</sub>O<sub>11</sub> | Low |
| M6 | -3.8 | 0.789 | 14.1 | +(O) | C<sub>33</sub>H<sub>57</sub>N<sub>3</sub>O<sub>10</sub> | Low |
| M7 | -3.7 | 0.800 | 14.3 | +(O) | C<sub>33</sub>H<sub>57</sub>N<sub>3</sub>O<sub>10</sub> | Low |
| M8 | -3.6 | 0.804 | 14.3 | -(H<sub>2</sub>) +(O<sub>2</sub>) | C<sub>33</sub>H<sub>55</sub>N<sub>3</sub>O<sub>11</sub> | Low |
| M9 | -3.2 | 0.814 | 14.5 | +(O) | C<sub>33</sub>H<sub>57</sub>N<sub>3</sub>O<sub>10</sub> | High |
| M10 | -3.0 | 0.832 | 14.8 | +(O) | C<sub>33</sub>H<sub>57</sub>N<sub>3</sub>O<sub>10</sub> | High |
| M11 | -2.7 | 0.848 | 15.1 | +(O) | C<sub>33</sub>H<sub>57</sub>N<sub>3</sub>O<sub>10</sub> | Low |
| M12 | -2.2 | 0.878 | 15.7 | -(C<sub>2</sub>H<sub>4</sub>) | C<sub>31</sub>H<sub>53</sub>N<sub>3</sub>O<sub>9</sub> | Low |
| M13 | -1.0 | 0.943 | 16.8 | -(CH<sub>2</sub>) | C<sub>32</sub>H<sub>55</sub>N<sub>3</sub>O<sub>9</sub> | High |
| Parent | 0.0 | 1.000 | 17.8 | None | C<sub>33</sub>H<sub>57</sub>N<sub>3</sub>O<sub>9</sub> | Low |
| **ENNB1** |  |  |  |  |  |  |
| M1 | -6.3 | 0.666 | 12.5 | -(H<sub>2</sub>) +(O<sub>2</sub>) | C<sub>34</sub>H<sub>57</sub>N<sub>3</sub>O<sub>11</sub> | Low |
| M2 | -5.4 | 0.710 | 13.3 | -(H<sub>2</sub>) +(O<sub>2</sub>) | C<sub>34</sub>H<sub>57</sub>N<sub>3</sub>O<sub>11</sub> | Low |
| M3 | -4.7 | 0.748 | 14.0 | -(H<sub>2</sub>) +(O<sub>2</sub>) | C<sub>34</sub>H<sub>57</sub>N<sub>3</sub>O<sub>11</sub> | Low |
| M4 | -4.7 | 0.752 | 14.1 | +(O) | C<sub>34</sub>H<sub>59</sub>N<sub>3</sub>O<sub>10</sub> | Low |
| M5 | -4.4 | 0.766 | 14.3 | -(H<sub>2</sub>) +(O<sub>2</sub>) | C<sub>34</sub>H<sub>57</sub>N<sub>3</sub>O<sub>11</sub> | Low |
| M6 | -4.0 | 0.788 | 14.8 | -(H<sub>2</sub>) +(O<sub>2</sub>) | C<sub>34</sub>H<sub>57</sub>N<sub>3</sub>O<sub>11</sub> | High |
| M7 | -3.8 | 0.800 | 15.0 | -(H<sub>2</sub>) +(O<sub>2</sub>) | C<sub>34</sub>H<sub>57</sub>N<sub>3</sub>O<sub>11</sub> | Low |
| M8 | -3.6 | 0.808 | 15.1 | +(O) | C<sub>34</sub>H<sub>59</sub>N<sub>3</sub>O<sub>10</sub> | High |
| M9 | -3.5 | 0.816 | 15.3 | +(O) | C<sub>34</sub>H<sub>59</sub>N<sub>3</sub>O<sub>10</sub> | High |
| M10 | -3.0 | 0.842 | 15.8 | -(H<sub>2</sub>) +(O) | C<sub>34</sub>H<sub>57</sub>N<sub>3</sub>O<sub>10</sub> | High |
| M11 | -2.4 | 0.870 | 16.3 | -(H<sub>2</sub>) +(O) | C<sub>34</sub>H<sub>57</sub>N<sub>3</sub>O<sub>10</sub> | High |
| M12 | -1.9 | 0.897 | 16.8 | -(C<sub>2</sub>H<sub>4</sub>) | C<sub>32</sub>H<sub>55</sub>N<sub>3</sub>O<sub>9</sub> | Low |
| M13 | -0.9 | 0.954 | 17.9 | -(CH<sub>2</sub>) | C<sub>33</sub>H<sub>57</sub>N<sub>3</sub>O<sub>9</sub> | Low |
| Parent | 0.0 | 1.000 | 18.7 | None | C<sub>34</sub>H<sub>59</sub>N<sub>3</sub>O<sub>9</sub> | High |

\*Δ retention time difference to peak of ENNB; \*\*Relative retention time (RT<sub>metabolite</sub>/RT<sub>ENNB</sub>).

Identification confidences varied between metabolites, of which some were low-abundant and had to be determined based on the sum of their sodiated and ammoniated ions. Major metabolites were hydroxylated (ENNB M9 and M10; ENNB1 M8 and M9) or carboxylated (ENNB1 M6, M10 and M11) (**Table 3**; **Supplementary Fig. 3**). The metabolites were detected in both compartments of the transwell system and in the cells (**Fig. 6**), showing that also the metabolites were transferred through the epithelial layer. The distribution into the compartments varied between the individual metabolites; e.g., whereas ENNB M3 remained apically and in the cells, ENNB M6 was almost completely transported into the BL compartment. Overall, hydroxylated metabolites were transported to a higher proportion than carboxylated or demethylated metabolites.

**Fig. 6.**
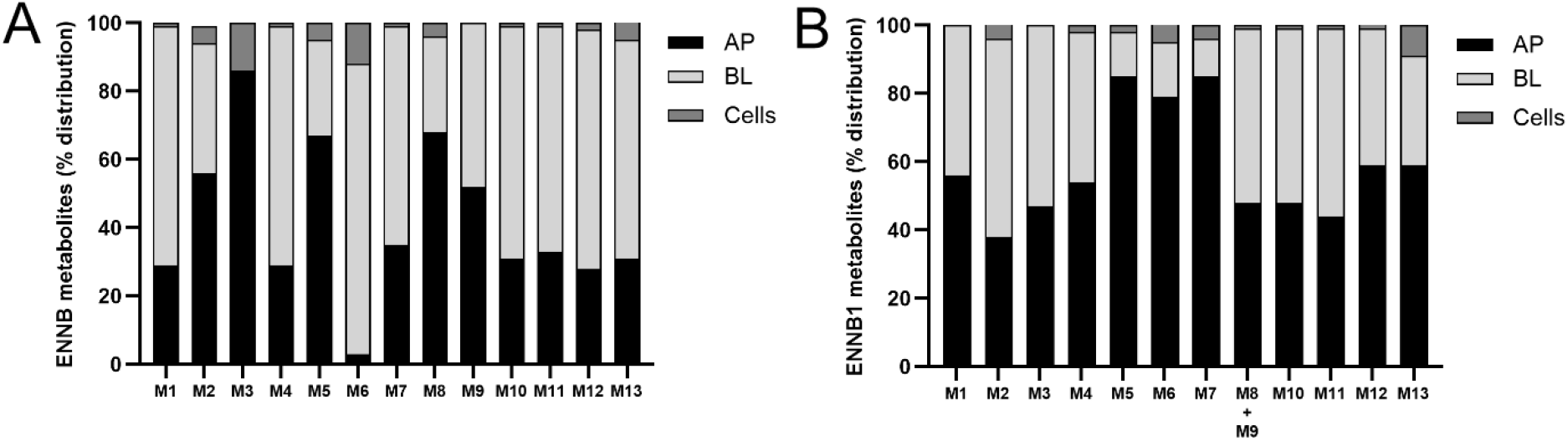
Distribution of **A**) ENNB, and **B**) ENNB1 metabolites in the compartments of the EpiIntestinal^TM^ tissues transwell system after apical application of 5 µM toxin and incubation for 4 h. Metabolites were identified by comparison to HLM results (**Supplementary** Fig. 3). Metabolite levels (of n=2 experiments) were determined semi-quantitatively based on the sodiated and ammoniated ions and used to calculate the distribution in relation to the total amount of each respective metabolite across all compartments.

The results of these experiments did, for the first time, not only confirm the ability of the primary human enterocytes in the EpiIntestinal^TM^ tissues to efficiently metabolise ENNB and ENNB1 to metabolites known from hepatic biotransformation studies but also showed that the model allowed determining the transmembrane transferability of the individual metabolites, which is relevant for the assessment of connected risks to human health. Thus, EpiIntestinal^TM^ tissues appeared to be a suitable model for determining xenobiotic biotransformation processes in enterocytes.

#### 3.3.3 Influence of ENNB on E2 metabolism

Several studies have shown that the ENNs and BEA can affect the biotransformation and Pg-p-dependent transport of other mycotoxins and physiological metabolites (Ivanova et al. 2019; Kalayou et al. 2015; Lori et al. 2026). Findings that they could interfere with steroidogenesis and have endocrine-disrupting effects are particularly concerning (Park and Lee 2021). The interaction between the ENNs and BEA and co-occurring compounds could be direct by competition for transporters or receptors, or indirect, e.g., by interaction with metabolic enzymes, which was investigated in the present study. When the sex hormone E2 was metabolised in the presence of different ENNB concentrations in HLM, the formation rates of its two main metabolites estrone (E1) and 2-hydroxy-estradiol (2-OH-E2) were increased (**Fig. 7**). Compared to the E2 control incubation, the addition of ENNB at concentrations 0.5 to 5 µM led to a 1.6-to 2.1-fold increase of E1 already after 15 min. The 2-OH-E2 levels also seemed to increase with the addition of ENNB; however, the increase was not significant. For all ENNB initial concentrations, the metabolite formation increased after starting the incubation, reached a plateau between 15 and 30 min, and slightly decreased at 60 min (data not shown).

**Fig. 7.**
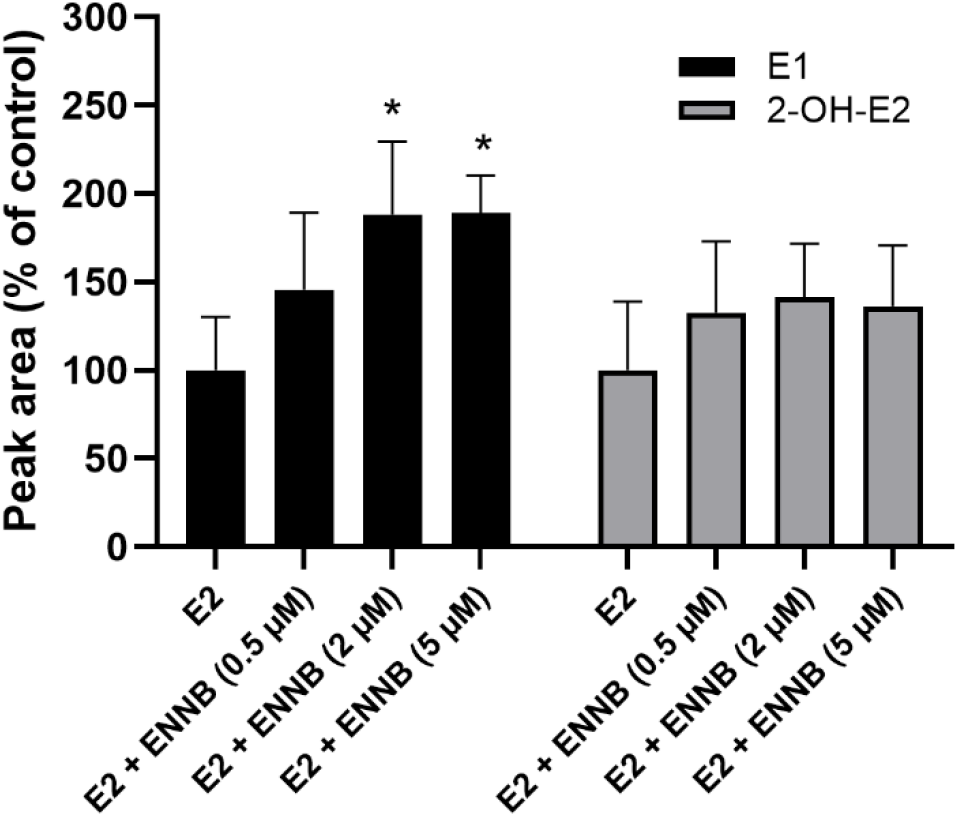
Impact of ENNB on the formation of the major E2 metabolites E1 and 2-OH-E2 in HLM (n=4). Initial E2 concentration 5 µM; metabolite formation (shown as ratios of HRMS peak areas compared to control E2) determined after incubation with different ENNB levels (0 to 5 µM) for 15 min. Significant differences (p< 0.05) compared to ctrl (0 µM) are indicated by an asterisk (*). Two-way ANOVA with Dunnett’s post hoc test.

The biotransformation pathway of E2 and the enzymes involved are well-established in humans (Tsuchiya et al. 2005). E1 is mostly formed through dehydrogenation catalysed by 17-beta-hydroxysteroid dehydrogenase (17β-HSD); however, CYP2C8, CYP2C9 and CYP2C19 can also drive the reaction. The hydroxylation of E2 to 2-OH-E2 is performed by CYP3A4 or CYP1A1/A2. The two major metabolites are subsequently metabolised to further products. Surprisingly, adding ENNB to E2 incubations in HLM increased metabolite formation, especially E1, the opposite of the enzyme inhibition usually observed for other anthropogenic and natural xenobiotics (Mallia et al. 2020; Usmani et al. 2006). It thus appeared that ENNB does not simply inhibit the respective CYP enzymes, but that the interaction is more complex. When the impact of ENNB1 on the CYP3A4-catalysed metabolism of midazolam (MDZ) previously was assessed in HLM, the elimination kinetics of the parent compound was slowed down significantly (Ivanova et al., 2019). However, analysis of the main MDZ metabolites showed that the formation of hydroxylated 4-OH-MDZ was increased by factor 4 after 30 min incubation. This accumulation of first-step metabolites of E2 and MDZ could indicate that ENNB and ENNB1 also inhibit metabolism enzymes further down the biotransformation pathway, e.g. the 16α-hydroxylation of E1, O-methylation of 2-OH-E2 or di-hydroxylation of 4-OH-MDZ. E1 has considerable oestrogenic potency, about one-third that of E2, so a significant increase in blood levels from dietary exposure to ENNs might disturb hormone balance and should be included in the assessment of potential risks.

In conclusion, the present study has demonstrated the applicability of human primary EpiIntestinal™ tissues to investigate the cytotoxicity, immunomodulation, transepithelial transport and intestinal biotransformation of xenobiotics such as the ENNs and BEA. The cell model could thus be used as an *in vitro* model for evaluating effects on the human intestine from dietary exposure to natural toxins and chemicals. Additionally, the immunotoxic potentials and transport properties of the ENNs and BEA were determined in a Caco-2/TH29-MTX-E12 co-culture, indicating a dependency between effect size and the molecular lipophilicity. Finally, the interaction of ENNB with biotransformation enzymes involved in E2 metabolism was evidenced in HLM, pointing at a possible interference with hormone homeostasis.

## Supporting information

Supplementary_Solhaug et al

## Acknowledgement

This work was carried out within the framework of the European Partnership for the Assessment of Risks from Chemicals (PARC) and has received funding from the European Union’s Horizon Europe research and innovation programme under Grant Agreement No 101057014. Views and opinions expressed are, however, those of the author(s) only and do not necessarily reflect those of the European Union or the Health and Digital Executive Agency. Neither the European Union nor the granting authority can be held responsible for them. The work was co-funded by the Norwegian Veterinary Institute (NVI). The authors would like to thank Brit Saure (NVI) for preparing and staining the EpiIntestinal^TM^FT tissues. The graphical abstract was created in BioRender. Solhaug, A. (2026) https://BioRender.com/t3sdobt

## Declaration of generative AI and AI-assisted technologies in the manuscript preparation process

During the preparation of this work, the authors used Grammarly pro (v1.2.305.1980) and Copilot (bizchat.20260701.51.17) to refine the language and improve the readability of the manuscript. After using these tools, the authors reviewed and edited the content as needed and take full responsibility for the content of the published article.

