## Supplementary_Solhaug et al for "Evaluation of immunotoxic effects and metabolism of enniatins and beauvericin using *in vitro* models of the human intestinal epithelium"

[*]Corresponding author:

**Supplementary Figure 1**

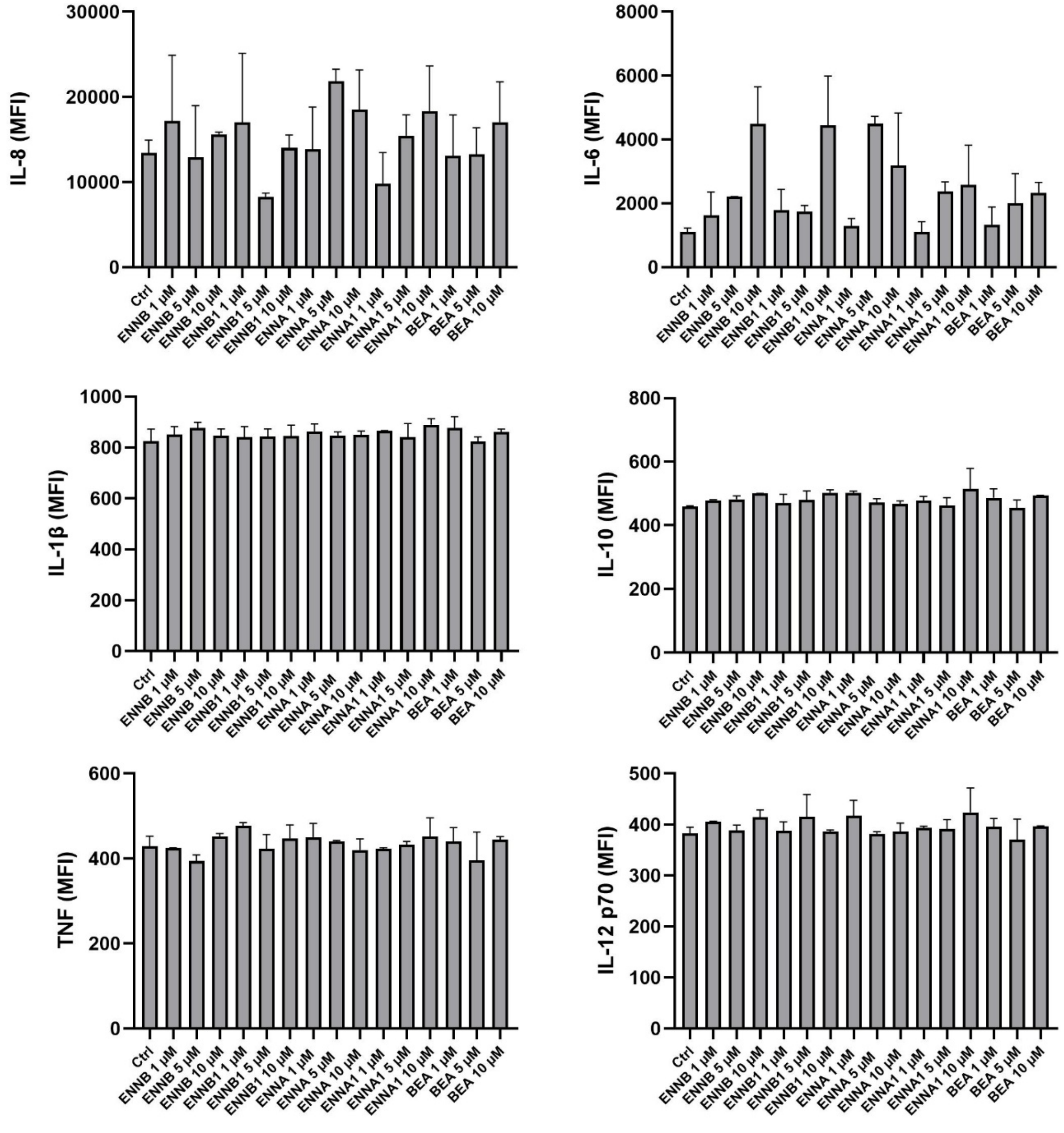


**Fig. 1:** Simultaneous measurement of IL-8, IL-6, TNFα, IL-1β, IL-10, and IL-12p70 secretion in EpiIntestinal™ tissues after exposure to ENNB, ENNB1, ENNA, ENNA1 and BEA (1, 5, 10 µM) for 24 h using the cytometric bead array (CBA). Data are expressed as mean + SD of two independent incubations. MFI = median fluorescent intensity, relative concentration.

**Supplementary Figure 2**

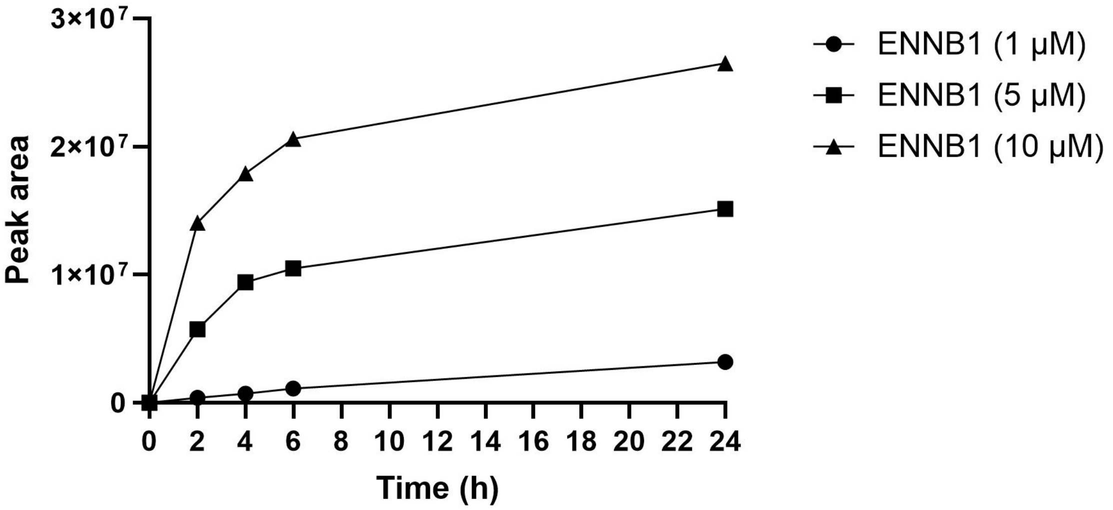


**Fig. 2**. Transport kinetics of ENNB1 in differentiated Caco-2 cells. Measured HRMS peak areas of ENNB1 in the BL compartment after AP exposure. One replicate performed for each concentration, of which aliquots were drawn after 0, 2, 4, 6 and 24 h of incubation.

**Supplementary Figure 3**

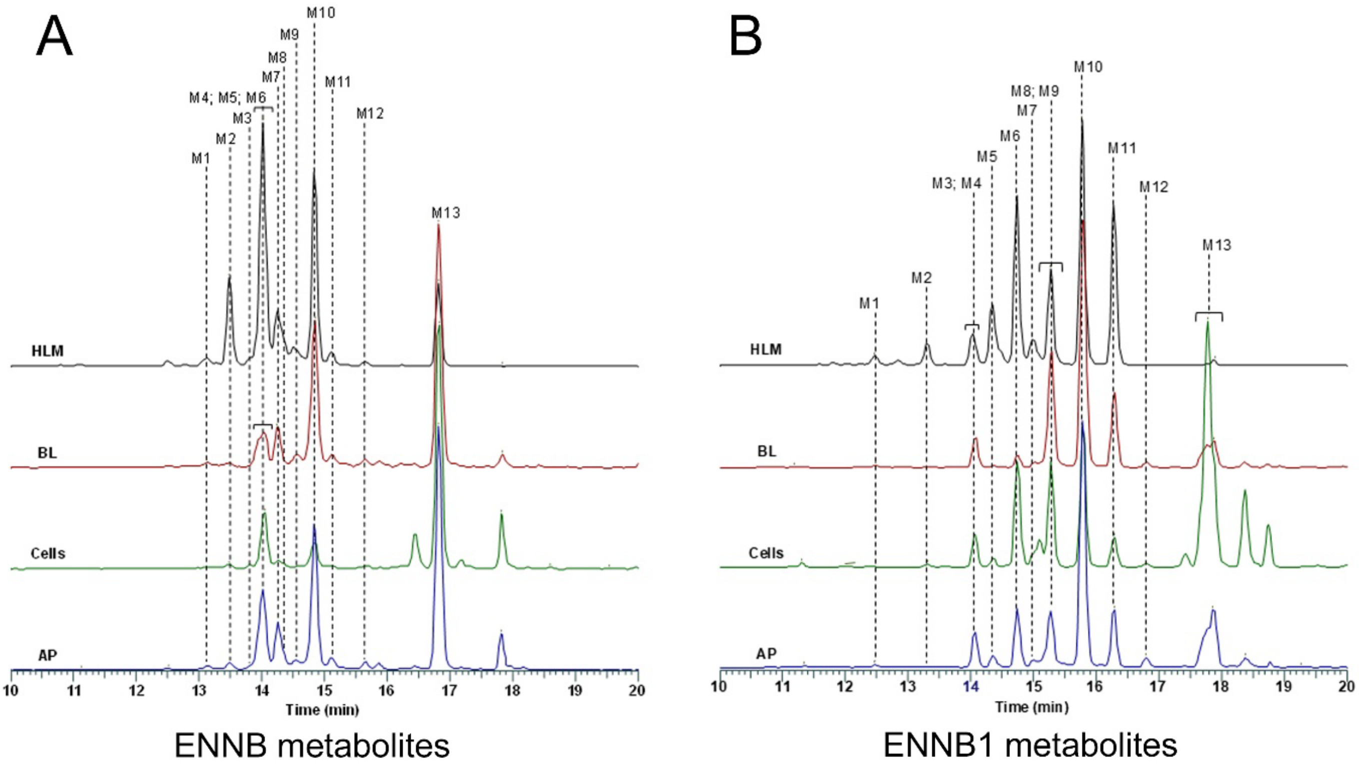


**Fig. 3. A)** ENNB and **B)** ENNB1 metabolites (M1 to M13; **Table 4**) determined in HLM, and the AP, BL and cell fractions of EpiIntestinal$^{TM}$ tissues. Each spectrum is normalised to the intensity of its most intense peak.

**Supplementary Table 1**

Validation parameters of the multi-toxin LC-HRMS analysis. Precision and recoveries of matrix-assisted calibration curves (Caco-2 cells or EpiIntestinal™ tissues).

| | **Caco-2** | | | | | **EpiIntestinal™ tissues** | | | | |
|---|---|---|---|---|---|---|---|---|---|---|
| | **conc (spike) µg/l** | **mean (n=6) µg/L** | **SD µg/L** | **intermediate precision RSD %** | **recovery %** | **conc (spike) µg/l** | **mean (n=2) µg/L** | **SD µg/L** | **intermediate precision RSD %** | **recovery %** |
| **ENNB** | 1 | 1.2 | 0.390 | 32 | 123 | 1 | 5.9 | - | - | 588 |
| | 5 | 5.1 | 0.376 | 7.4 | 102 | 5 | 9.0 | - | - | 180 |
| | 10 | 10.1 | 0.582 | 5.8 | 101 | 10 | 12.8 | 0.111 | 0.9 | 128 |
| | 50 | 50.6 | 3.190 | 6.3 | 101 | 50 | 44.9 | 1.037 | 2.3 | 90 |
| | 100 | 102.2 | 6.913 | 6.8 | 102 | 100 | 90.5 | 1.073 | 1.2 | 90 |
| | 250 | 242.3 | 15.09 | 6.2 | 97 | 250 | 255.1 | 20.25 | 7.9 | 102 |
| | 500 | 519.0 | 32.13 | 6.2 | 104 | 500 | 509.9 | 35.67 | 7.0 | 102 |
| **ENNB1** | | | | | | | | | | |
| | 1 | 0.8 | 0.223 | 28 | 80 | 1 | 2.3 | - | - | 226 |
| | 5 | 4.8 | 0.246 | 5.1 | 95 | 5 | 6.0 | 0.171 | 2.8 | 120 |
| | 10 | 9.8 | 0.485 | 4.9 | 98 | 10 | 10.6 | 0.062 | 0.6 | 106 |
| | 50 | 50.6 | 3.419 | 6.8 | 101 | 50 | 48.7 | 1.085 | 2.2 | 97 |
| | 100 | 101.2 | 6.618 | 6.5 | 101 | 100 | 98.5 | 2.566 | 2.6 | 98 |
| | 250 | 239.1 | 13.62 | 5.7 | 96 | 250 | 250.6 | 9.752 | 3.9 | 100 |
| | 500 | 511.3 | 31.58 | 6.2 | 102 | 500 | 502.2 | 9.172 | 1.8 | 100 |
| **ENNA** | | | | | | | | | | |
| | 1 | 0.6 | 0.597 | 108 | 55 | 1 | 4.6 | - | - | 462 |
| | 5 | 4.5 | 0.509 | 11 | 91 | 5 | 7.3 | 0.081 | 1.1 | 146 |
| | 10 | 10.1 | 0.658 | 6.5 | 101 | 10 | 11.2 | 0.098 | 0.9 | 112 |
| | 50 | 52.4 | 3.597 | 6.9 | 105 | 50 | 48.3 | 0.026 | 0.1 | 97 |
| | 100 | 103.8 | 6.330 | 6.1 | 104 | 100 | 97.6 | 1.308 | 1.3 | 98 |
| | 250 | 236.8 | 10.70 | 4.5 | 95 | 250 | 243.4 | 4.681 | 1.9 | 97 |
| | 500 | 509.2 | 21.70 | 4.3 | 102 | 500 | 505.4 | 1.125 | 0.2 | 101 |
| **ENNA1** | | | | | | | | | | |
| | 1 | 0.7 | 0.465 | 71 | 65 | 1 | 0.6 | - | - | 63 |
| | 5 | 4.7 | 0.351 | 7.5 | 94 | 5 | 4.3 | - | - | 86 |
| | 10 | 9.8 | 0.587 | 6.0 | 98 | 10 | 8.9 | - | - | 89 |
| | 50 | 51.8 | 3.905 | 7.5 | 104 | 50 | 47.2 | 0.679 | 1.4 | 94 |
| | 100 | 102.5 | 6.564 | 6.4 | 102 | 100 | 98.9 | 1.578 | 1.6 | 99 |
| | 250 | 237.8 | 12.34 | 5.2 | 95 | 250 | 260.0 | 8.406 | 3.2 | 104 |
| | 500 | 509.2 | 27.69 | 5.4 | 102 | 500 | 496.6 | 17.32 | 3.5 | 99 |
| **BEA** | | (n=2) | | | | | | | | |
| | 1 | 13.5 | - | - | 1352 | 1 | 6.2 | - | - | 620 |
| | 5 | - | - | - | - | 5 | 9.3 | 0.080 | 0.9 | 186 |
| | 10 | 16.9 | 0.189 | 1.1 | 169 | 10 | 13.7 | 0.019 | 0.1 | 137 |
| | 50 | 43.0 | 4.788 | 11 | 86 | 50 | 46.2 | 0.885 | 1.9 | 92 |
| | 100 | 92.4 | 0.206 | 0.2 | 92 | 100 | 93.6 | 0.915 | 1.0 | 94 |
| | 250 | 242.6 | 0.000 | 0.0 | 97 | 250 | 250.1 | 6.836 | 2.7 | 100 |
| | 500 | 512.7 | 24.92 | 4.9 | 103 | 500 | 513.2 | 8.614 | 1.7 | 103 |

SD: standard deviation; RSD%: relative standard deviation;

**Supplementary Table 2**

Total toxin recoveries across all compartments (AP, BL, cells) in Caco-2 cells and EpiIntestinal™ tissues in transwell inserts after apical application of 5 µM ENNs and BEA, with or without the inhibitor PSC833, and incubation for 4 h. ENNs and BEA concentrations were determined by LC-HRMS analysis, amounts calculated under consideration of the compartment volumes, summed and compared to the amount added at 0 h. Values are shown as mean ± SD from 3-4 independent experiments (Caco-2) or as mean ± SD from 2 independent incubations (EpiIntestinal™ tissues). Significant differences ($p < 0.05$) as compared to ENNs/BEA without inhibitor are indicated with an asterisk (*) (unpaired t-test).

| Total toxin recoveries (%) | | |
|---|---|---|
| **Toxin ± PSC833** | **Caco-2** | **EpiIntestinal™ tissues** |
| ENNB | 92 ± 15 | 93 ± 2 |
| ENNB + PSC833 | 98 ± 28 | 77 ± 1* |
| ENNB1 | 114 ± 6 | 106 ± 6 |
| ENNB1 + PSC833 | 104 ± 9 | 76 ± 4* |
| ENNA | 94 ± 20 | 105 ± 18 |
| ENNA + PSC833 | 96 ± 20 | 97 ± 2 |
| ENNA1 | 89 ± 21 | 80 ± 2 |
| ENNA1 + PSC833 | 89 ± 21 | 70 ± 3 |
| BEA | 80 ± 5 | 83 ± 1 |
| BEA + PSC833 | 79 ± 17 | 55 ± 2* |

### Supplementary Table 3

ENNB metabolites detected across the compartments of EpiIntestinal™ tissues in transwell inserts after incubation with 5 µM ENNB for 4h. Combined results of two experiments.

| ID | Name | ΔRT | $RT_{ratio}$ | RT | Change | Formula | $[M+NH_4]^+$ | | | | | | | | $[M+Na]^+$ | | | | | | | | Confid. |
|---|---|---|---|---|---|---|---|---|---|---|---|---|---|---|---|---|---|---|---|---|---|---|---|
| | | | | | | | Observed mass (ppm) | | | Theor. mass (ppm) | Diff. to theor. mass (ppm) | | | | Observed mass (ppm) | | | Theor. mass (ppm) | Diff. to theor. mass (ppm) | | | | |
| | | | | | | | AP | BL | Cells | | AP | BL | Cells | | AP | BL | Cells | | AP | BL | Cells | | |
| **M1** | Enniatin B_+O/-$CH_2$_Δ -4.8 | -4.7 | 0.737 | 13.1 | -($CH_2$) +(O) | $C_{32}H_{55}N_3O_{10}$ | 659.4236 | 659.4226 | 659.4228 | 659.4226 | 1.5 | 0.06 | 0.3 | | NF | NF | NF | 664.3780 | NF | NF | NF | | low |
| **M2** | Enniatin B_+2O/-2H_Δ -4.5 | -4.4 | 0.756 | 13.5 | -(2H) +(2O) | $C_{33}H_{55}N_3O_{11}$ | 687.4181 | 687.4175 | 687.4178 | 687.4175 | 0.9 | -0.01 | 0.5 | | 692.3723 | NF | NF | 692.3729 | -0.8 | NF | NF | | low |
| **M3** | Enniatin B_+2O/-2H_Δ -4.1 | -4.0 | 0.774 | 13.8 | -(2H) +(2O) | $C_{33}H_{55}N_3O_{11}$ | 687.4182 | NF | 687.4175 | 687.4175 | 1.1 | NF | -0.01 | | NF | NF | NF | 692.3729 | NF | NF | NF | | low |
| **M4** | Enniatin B_+O/-$CH_2$_Δ -3.9 | -3.9 | 0.782 | 13.9 | -($CH_2$) +(O) | $C_{32}H_{55}N_3O_{10}$ | 659.4238 | 659.4236 | 659.4227 | 659.4226 | 1.9 | 1.5 | 0.2 | | 664.3788 | 664.3788 | NF | 664.3780 | 1.3 | 1.3 | NF | | low |
| **M5** | Enniatin B_+2O/-2H_Δ -3.9 | -3.8 | 0.786 | 14.0 | -(H2) +(O2) | $C_{33}H_{55}N_3O_{11}$ | 687.4183 | 687.4184 | 687.4178 | 687.4175 | 1.2 | 1.3 | 0.4 | | 692.3737 | NF | 692.3730 | 692.3729 | 1.1 | NF | 0.2 | | low |
| **M6** | Enniatin B_O_Δ -3.8 | -3.8 | 0.789 | 14.1 | +(O) | $C_{33}H_{57}N_3O_{10}$ | 673.4388 | 673.4389 | 673.4388 | 673.4382 | 0.9 | 1.0 | 0.8 | | NF | NF | NF | 678.3936 | NF | NF | NF | | low |
| **M7** | Enniatin B_O_Δ -3.7 | -3.6 | 0.800 | 14.3 | +(O) | $C_{33}H_{57}N_3O_{10}$ | 673.4390 | 673.4391 | 673.4385 | 673.4382 | 1.1 | 1.4 | 0.5 | | 678.3942 | 678.3941 | NF | 678.3936 | 0.8 | 0.7 | NF | | low |
| **M8** | Enniatin B_+2O/-2H_Δ -3.6 | -3.5 | 0.804 | 14.3 | -(2H) +(2O) | $C_{33}H_{55}N_3O_{11}$ | 687.4180 | 687.4178 | 687.4164 | 687.4175 | 0.8 | 0.5 | -1.5 | | 692.3724 | NF | NF | 692.3729 | 0.7 | NF | NF | | low |
| **M9** | Enniatin B_O_Δ -3.2 | -3.3 | 0.814 | 14.5 | +(O) | $C_{33}H_{57}N_3O_{10}$ | 673.4386 | 673.4387 | NF | 673.4382 | 0.6 | 0.7 | NF | | 678.3940 | 678.3942 | NF | 678.3936 | 0.6 | 0.9 | NF | | high |
| **M10** | Enniatin B_O_Δ -3.0 | -3.0 | 0.832 | 14.8 | +(O) | $C_{33}H_{57}N_3O_{10}$ | 673.4393 | 673.4393 | 673.4389 | 673.4382 | 1.7 | 1.6 | 1.0 | | 678.3945 | 678.3944 | 678.3936 | 678.3936 | 1.4 | 1.2 | -0.1 | | high |
| **M11** | Enniatin B_O_Δ -2.7 | -2.7 | 0.848 | 15.1 | +(O) | $C_{33}H_{57}N_3O_{10}$ | 673.4391 | 673.4387 | NF | 673.4382 | 1.4 | 0.7 | NF | | 678.3945 | NF | NF | 678.3936 | 1.3 | NF | NF | | low |
| **M12** | Enniatin B_-$C_2H_4$_Δ -2.2 | -2.2 | 0.878 | 15.7 | -($C_2H_4$) | $C_{31}H_{53}N_3O_9$ | 629.4126 | 629.4119 | 629.4113 | 629.4120 | 0.9 | -0.1 | -1.1 | | NF | NF | NF | 634.3674 | NF | NF | NF | | low |
| **M13** | Enniatin B_-$CH_2$_Δ -1.0 | -1.0 | 0.943 | 16.8 | -($CH_2$) | $C_{32}H_{55}N_3O_9$ | 643.4291 | 643.4290 | 643.4286 | 643.4277 | 2.2 | 2.0 | 1.5 | | 648.3842 | 648.3842 | 648.3837 | 648.3831 | 1.7 | 1.8 | 1.0 | | high |
| **ENNB** | Enniatin B_Δ 0 | 0.0 | 1.000 | 17.8 | None | $C_{33}H_{57}N_3O_9$ | 657.4442 | 657.4437 | 657.4438 | 657.4433 | 1.4 | 0.6 | 0.7 | | 662.3987 | 662.3984 | 662.3986 | 662.3987 | 0.1 | -0.4 | -0.2 | | low |

RT: retention time; AP: apical compartment; BL: basolateral compartment; NF: not found

### Supplementary Table 4

ENNB1 metabolites detected across the compartments of EpiIntestinal™ tissues in transwell inserts after incubation with 5 µM ENNB for 4h. Combined results of two experiments.

| ID | Name | ΔRT | $RT_{ratio}$ | RT | Change | Formula | $[M+NH_4]^+$ Observed mass (ppm) | | | $[M+NH_4]^+$ Theor. mass (ppm) | $[M+NH_4]^+$ Diff. to theor. mass (ppm) | | | $[M+Na]^+$ Observed mass (ppm) | | | $[M+Na]^+$ Theor. mass (ppm) | $[M+Na]^+$ Diff. to theor. mass (ppm) | | | Confid. |
|---|---|---|---|---|---|---|---|---|---|---|---|---|---|---|---|---|---|---|---|---|---|
| | | | | | | | AP | BL | Cells | | AP | BL | Cells | AP | BL | Cells | | AP | BL | Cells | |
| **M1** | Enniatin B1_+2O/-2H_Δ -6.3 | -6.3 | 0.666 | 12.5 | -(2H) +(2O) | $C_{34}H_{57}N_3O_{11}$ | 701.4327 | 701.4314 | NF | 701.4331 | -0.7 | -2.5 | NF | 706.3879 | 706.3865 | NF | 706.3885 | -0.9 | -2.8 | NF | low |
| **M2** | Enniatin B1_+2O/-2H_Δ -5.5 | -5.4 | 0.710 | 13.3 | -(2H) +(2O) | $C_{34}H_{57}N_3O_{11}$ | 701.4322 | 701.4316 | 701.4307 | 701.4331 | -1.4 | -2.2 | -3.5 | 706.3871 | 706.3874 | 706.3870 | 706.3885 | -2.1 | -1.6 | -2.2 | low |
| **M3** | Enniatin B1_+2O/-2H_Δ -4.7 | -4.7 | 0.748 | 14.0 | -(2H) +(2O) | $C_{34}H_{57}N_3O_{11}$ | 701.4323 | 701.4327 | NF | 701.4331 | -1.2 | -0.6 | NF | 706.3892 | NF | NF | 706.3885 | 0.9 | NF | NF | low |
| **M4** | Enniatin B1_O_Δ -4.7 | -4.7 | 0.752 | 14.1 | +(O) | $C_{34}H_{59}N_3O_{10}$ | 687.4532 | 687.4533 | 687.4537 | 687.4539 | -1.0 | -0.8 | -0.3 | 692.4085 | 692.4085 | 692.4081 | 692.4093 | -1.1 | -1.1 | -1.6 | low |
| **M5** | Enniatin B1_+2O/-2H_Δ -4.5 | -4.4 | 0.766 | 14.3 | -(2H) +(2O) | $C_{34}H_{57}N_3O_{11}$ | 701.4326 | 701.4319 | 701.4324 | 701.4331 | -0.8 | -1.7 | -1.1 | 706.3879 | 706.3870 | 706.3900 | 706.3885 | -0.8 | -2.1 | 2.0 | low |
| **M6** | Enniatin B1_+2O/-2H_Δ -4.0 | -4.0 | 0.788 | 14.8 | -(2H) +(2O) | $C_{34}H_{57}N_3O_{11}$ | 701.4328 | 701.4323 | 701.4326 | 701.4331 | -0.5 | -1.3 | -0.8 | 706.3881 | 706.3871 | 706.3881 | 706.3885 | -0.7 | -2.0 | -0.6 | high |
| **M7** | Enniatin B1_+2O/-2H_Δ -3.8 | -3.8 | 0.800 | 15.0 | -(2H) +(2O) | $C_{34}H_{57}N_3O_{11}$ | 701.4324 | 701.4323 | 701.4324 | 701.4331 | -1.0 | -1.2 | -1.0 | 706.3877 | NF | 706.3902 | 706.3885 | -1.2 | NF | 2.4 | low |
| **M8** | Enniatin B1_O_Δ -3.6 | -3.6 | 0.808 | 15.1 | +(O) | $C_{34}H_{59}N_3O_{10}$ | 687.4536 | 687.4534 | 687.4535 | 687.4539 | -0.4 | -0.6 | -0.6 | 692.4088 | 692.4087 | 692.4086 | 692.4093 | -0.7 | -0.8 | -1.0 | high |
| **M9** | Enniatin B1_O_Δ -3.5 | -3.5 | 0.816 | 15.3 | +(O) | $C_{34}H_{59}N_3O_{10}$ | 687.4538 | 687.4535 | 687.4535 | 687.4539 | -0.1 | -0.6 | -0.6 | 692.4090 | 692.4088 | 692.4087 | 692.4093 | -0.4 | -0.7 | -0.8 | high |
| **M10** | Enniatin B1_+O/-2H_Δ -3.0 | -3.0 | 0.842 | 15.8 | -(2H) +(O) | $C_{34}H_{57}N_3O_{10}$ | 685.4380 | 685.4380 | 685.4376 | 685.4382 | -0.3 | -0.3 | -1.0 | 690.3930 | 690.3933 | 690.3930 | 690.3936 | -0.9 | -0.4 | -0.9 | high |
| **M11** | Enniatin B1_+O/-2H_Δ -2.4 | -2.4 | 0.870 | 16.3 | -(2H) +(O) | $C_{34}H_{57}N_3O_{10}$ | 685.4380 | 685.4377 | 685.4376 | 685.4382 | -0.3 | -0.8 | -1.0 | 690.3932 | 690.3931 | 690.3922 | 690.3936 | -0.6 | -0.7 | -2.1 | high |
| **M12** | Enniatin B1_-C2H4_Δ -1.9 | -1.9 | 0.897 | 16.8 | $-(C_2H_4)$ | $C_{32}H_{55}N_3O_9$ | 643.4272 | 643.4269 | 643.4271 | 643.4277 | -0.6 | -1.2 | -0.8 | 648.3823 | 648.3817 | NF | 648.3831 | -1.1 | -2.1 | NF | low |
| **M13** | Enniatin B1_-CH2_Δ -0.9 | -0.9 | 0.954 | 17.9 | $-(CH_2)$ | $C_{33}H_{57}N_3O_9$ | 657.4430 | 657.4431 | 657.4431 | 657.4433 | -0.5 | -0.4 | -0.3 | 662.3983 | 662.3987 | 662.3983 | 662.3987 | -0.6 | 0.1 | -0.7 | low |
| **ENNB1** | Enniatin B1_Δ -0 | 0.0 | 1.000 | 18.7 | None | $C_{34}H_{59}N_3O_9$ | 671.4583 | 671.4589 | 671.4583 | 671.4590 | -1.1 | -0.1 | -1.1 | 676.4128 | 676.4131 | 676.4128 | 676.4144 | -2.2 | -1.8 | -2.2 | high |

RT: retention time; AP: apical compartment; BL: basolateral compartment; NF: not found
